# Global O-GlcNAcome Identifies O-GlcNAcylated Proteins Regulating Oligodendrocyte Precursor Cells Differentiation

**DOI:** 10.64898/2026.03.02.709210

**Authors:** Xinyue Jiang, Chi Zhang, Pingping Li, Ke Ye, Yixuan Song, Mingliang Zhang

## Abstract

Oligodendrocyte precursor cells (OPCs) differentiate into myelinating oligodendrocytes to form the myelin sheaths essential for neural function, a process frequently compromised during aging and in diseases like multiple sclerosis. However, the regulatory landscape of myelination remains incompletely understood. Here, we perform a comprehensive proteomic analysis of the O-GlcNAcome in postnatal day 0 mouse OPCs, identifying 165 O-GlcNAcylation sites across 118 proteins. These proteins are mostly nuclear and characterized by single-site modifications. Comparative analysis identifies 74 novel sites and 22 proteins uniquely O-GlcNAcylated in OPCs. Molecular docking suggestes that O-GlcNAcylation might act as a dynamic regulator of OPC differentiation. Notably, global O-GlcNAcylation levels increase during the OPC-to-oligodendrocyte transition. By integrating our findings with multiple sclerosis-related transcriptomic and proteomic datasets, we identify candidate proteins linking O-GlcNAc dysregulation to demyelination. Among these, *Vimentin* (*Vim*) exhibits a pronounced age-dependent discrepancy between decreased protein levels and elevated mRNA expression during differentiation. We validate Threonine 35 and 63 as key O-GlcNAcylation sites on Vimentin. Functional assays confirm that blocking Vimentin O-GlcNAcylation enhance the OPC differentiation. Collectively, our study establishes the first O-GlcNAc landscape for OPCs and suggests O-GlcNAcylation as a pivotal regulator for OPC differentiation.

## Introduction

In the central nervous system (CNS), oligodendrocyte precursor cells (OPCs) differentiate into mature oligodendrocytes (OLs) to produce myelin sheaths. Myelin structures are essential for insulating neuronal axons, facilitating saltatory conduction, and providing metabolic support [1–4]. This developmental process represents a cornerstone of CNS plasticity and remyelination [5–8]. OPC differentiation depends on tightly coordinated gene expression, metabolic reprogramming, and cytoskeletal reorganization [9]. Consequently, dysregulation of this program could lead to severe pathologies, such as pediatric leukodystrophies and impaired myelin regeneration in adults [10]. In multiple sclerosis, a chronic autoimmune disorder characterized by demyelination, OPCs recruited to lesion sites often fail to differentiate into mature OLs, resulting in progressive axonal degeneration and disability [11–14]. Similarly, aging significantly reduces the regenerative capacity of OPCs, contributing to white matter loss and cognitive decline [15, 16]. Despite substantial progress in OPC biology, the molecular and cellular drivers of differentiation remain incompletely understood. While certain remyelination regulators, such as muscarinic receptor pathways, are increasingly well defined [17], the post-translational modifications (PTMs) governing these processes is still largely unexplored.

The transition from OPCs to OLs is a metabolically intensive process that requires the coordination of the integration of cellular nutrient status with lineage progression [18]. A central mediator of this nutrient sensing is O-linked β-N-acetylglucosaminylation (O-GlcNAcylation), a dynamic PTM involving the attachment of β-N-acetylglucosamine to serine or threonine residues. This modification is regulated by two primary enzymes, O-GlcNAc transferase (Ogt) and O-GlcNAcase (Oga) [19–21].

As a nutrient sensor, O-GlcNAcylation utilizes uridine diphosphate N-acetylglucosamine (UDP-GlcNAc) from the hexosamine biosynthetic pathway, converging glucose, amino acid, lipid, and nucleotide metabolism [22–24]. While this modification is abundant in the CNS and critical for neural development, synaptic function, and neurogenesis [25, 26], its dysregulation is frequently linked to neurodegenerative conditions such as Alzheimer’s and Parkinson’s diseases [27–30]. In the context of multiple sclerosis, reduced GlcNAc levels have been correlated with disease progression, whereas supplementation has been shown to promote OL differentiation [31, 32]. However, the specific O-GlcNAc-mediated mechanisms that drive OPC lineage progression remain to be elucidated.

Recent breakthroughs in mass spectrometry-based proteomics, supported by innovative chemical probes and chemoenzymatic labeling, have revolutionized the global characterization of the O-GlcNAcome [33]. Detailed mapping of these modifications has provided profound insights across various physiological contexts. For instance, landmark studies in murine synapses identified over 1,750 sites on proteins involved in vesicle trafficking and cytoskeletal organization [34]. Quantitative approaches have further enabled precise mapping of O-GlcNAcylation dynamics [35]. Specifically, quantitative proteomics in post-mortem human Alzheimer’s disease brain tissue, utilizing chemoenzymatic photocleavage enrichment and TMT labelling, revealed 131 O-GlcNAcylated peptides with significant disease-associated changes [36]. Normalization to total protein abundance confirmed these alterations reflected occupancy shifts rather than expression changes, with affected proteins enriched in synaptic pathways. In non-neural contexts, studies in mouse embryonic stem cells have demonstrated that elevated global O-GlcNAcylation is essential for maintaining pluripotency and must decrease to permit differentiation. This modification establishes a stemness-preserving network, as evidenced by the identification of 979 O-GlcNAcylated proteins that regulate the undifferentiated state [37]. These efforts have culminated in valuable resources, including the O-GlcNAcAtlas and the O-GlcNAc Database, which curate thousands of identified sites [38, 39]. Despite this progress in mapping the general O-GlcNAcome, the O-GlcNAcylation landscape of OPCs and how it dynamically shifts during lineage progression remains unexplored.

In this study, we characterized the O-GlcNAcome of postnatal day 0 (P0) mouse OPCs, mapping O-GlcNAcylated proteins and identified previously uncharacterized modification sites. These modified proteins are enriched in pathways related to cytoskeletal organization and neural development. By integrating our O-GlcNAcome dataset with the transcriptomic and proteomic profiles from pathological (multiple sclerosis) and physiological (aging) contexts, we elucidated the dynamic changes in O-GlcNAcylated proteins across disease and aging. This analysis further pinpointed candidate regulators, including *Vimentin*, whose expression levels vary temporally in these conditions. Functional validation confirmed that O-GlcNAcylation of Vimentin is crucial for OPC differentiation. Together, these results provide the first O-GlcNAcome profile for OPCs, offering a potential therapeutic strategy for myelin regeneration.

## Materials and Methods

### Animals

All animal experiments were conducted in accordance with the guidelines approved by the Institutional Animal Care and Use Committee of Shanghai Jiao Tong University School of Medicine (Approval Number JUMC2023-129-A). C57BL/6 mice were housed in a specific-pathogen-free (SPF) facility under a controlled 12-hour light/dark cycle, with ad libitum access to standard chow and water. Timed-pregnant females were used for the isolation of embryonic neural precursor cells, with the day of vaginal plug detection designated as embryonic day 0.5 (E0.5).

### Brain tissue collection and processing for immunostaining

For the *in vivo* analysis of O-GlcNAcylation, postnatal day 14 (P14), 2-month-old and 12-month-old mice were deeply anesthetized and transcardially perfused with ice-cold phosphate-buffered saline (PBS, pH 7.4) followed by 4% paraformaldehyde (PFA) in PBS for fixation. Brains were dissected, post-fixed in 4% PFA at 4°C overnight, and coronally sectioned at 30 µm using a vibrating microtome (Leica VT1200 S, Wetzlar, Germany). Sections were then carefully transferred into 24-well plates containing PBS and stored at 4°C for immunostaining.

### Primary neural progenitor cells isolation and culture

For *in vitro* differentiation and functional assays, primary neural progenitor cells (pNPCs) were isolated from the cerebral cortices of embryonic day 12.5 (E12.5) C57BL/6J mouse embryos under sterile conditions in a laminar flow hood. Brains were dissected in ice-cold PBS, and cerebral hemispheres were separated. Meninges were removed using fine forceps, and cortices were minced and dissociated using Trpzyme (BasalMedia, Shanghai, China) for 15–20 minutes at 37°C. Digestion was stopped with 10% fetal bovine serum (FBS; LONSERA, Suzhou, China) in Dulbecco’s Modified Eagle Medium (DMEM; BasalMedia). The cell suspension was filtered through a 40µm strainer, centrifuged at 1000 rpm for 5 minutes, and resuspended in NPC culture medium consisting of Neurobasal Medium (Gibco, Waltham, USA) and DMEM/F12 (Gibco) (1:1) supplemented with 1× N2 (Gibco), 1× B27 without VA (Gibco), 1× non-essential amino acids (NEAA) (Gibco), 1× GlutaMAX (Gibco), 0.075% Bovine serum albumin (BSA; YEASEN, Shanghai, China), 20 ng/ml mouse epidermal growth factor (mEGF; Peprotech, Rocky Hill, USA), 20 ng/ml basic fibroblast growth factor (bFGF; Peprotech), and 1× penicillin-streptomycin (PS; BasalMedia). Cells were cultured as neurospheres in uncoated 10-cm dishes for suspension culture in a humidified incubator at 37°C with 5% CO[, with partial medium replacement daily.

Neurospheres reaching ∼200 µm in diameter were collected in a 50-mL centrifuge tube and allowed to settle for 3 minutes. The supernatant and unsettled cells were discarded, and 2 mL pre-warmed TrypZyme was added to digest the neurospheres at 37°C for 5 minutes. Digestion was stopped with 10% FBS in DMEM, and cells were centrifuged at 1000 rpm for 3 min. The pellet was resuspended in 5 mL pre-warmed NPC medium. Consequently, the cells were counted and seeded at 3 x 10^4^ cells/cm^2^ in Matrigel (Corning, Corning, USA) coated well plates and cultured in fresh NPC medium at 37°C and 5% CO_2_.

### Differentiation of NPCs into oligodendrocyte precursor cells and oligodendrocytes

NPCs were seeded onto Matrigel-coated well plates at a density of 2.5 × 10^4^ cells/cm^2^ in OPC medium [DMEM/F12 basal medium supplemented with 1× N2, 1× B27 (without vitamin A), 1× GlutaMAX, 1× NEAA, 0.1 mM β-mercaptoethanol (Gibco), 200 ng/mL sonic hedgehog (shh; R&D Systems, Minneapolis, USA), 20 ng/mL bFGF, and 20 ng/mL platelet-derived growth factor-AA (PDGF-AA; Peprotech)]. The cells were maintained in OPC medium for 3 days, with the medium refreshed every other day. OPC medium was then replaced with OL medium [DMEM/F12 supplemented with 1× N2, 1× B27 (without vitamin A), 1× GlutaMAX, 1× NEAA, 0.1 mM β-mercaptoethanol, 200 ng/mL Shh, 100 nM LDN193189, 100 ng/m insulin-like growth factor 1 (IGF-1), 10 mM cyclic AMP, 10 ng/mL neurotrophin-3 (NT-3; Peprotech), and 40 ng/mL triiodothyronine (T3; MCE, Monmouth Junction, USA)] for 4–5 days of culture, with half of the culture medium replaced every other day. During this process, the cells were maintained in an incubator at a constant temperature of 37°C and at 5% CO_2_.

### Assessment of cell culture purity

To assess the purity of OPC and OL cultures derived from NPCs, cells were immunostained with lineage-specific markers. Purity was determined by calculating the percentage of platelet-derived growth factor receptor alpha (Pdgfrα) positive cells (for OPCs) or myelin basic protein (Mbp) positive cells (for OLs) relative to the total number of 4′,6[diamidino[2[phenylindole (DAPI)-stained nuclei. For primary OPCs isolated from P0 mice for proteomics, purity was assessed via flow cytometry using an anti-Pdgfrα antibody (Invitrogen). An isotype control antibody was used to establish the background signal.

### Functional analysis of O-GlcNAc cycling enzymes in NPCs

To investigate the effect of O-GlcNAc cycling on differentiation, primary NPCs were transfected with pCDNA3.1-*Oga*, pCDNA3.1-*Ogt* overexpression plasmids, or an empty vector control using the Neon Transfection System. Following transfection, cells were subjected to bleomycin selection for 3 days to enrich for transfected cells. The selected cells were then induced to differentiate into OPCs or mature OLs using the differentiation protocols described above. Differentiation efficiency was assessed by immunostaining for O4 and Mbp.

### Culture of CG4 cells

CG4 cells were maintained in DMEM-F12 medium supplied with N2, B27, GlutaMAX, PDGF-AA and mEGF as described. Differentiation was induced by differentiation medium supplemented with B27, GlutaMAX, 50mg/mL Transferrin (BasalMedia), 50 ng/mL ciliary neurotrophic factor (CNTF; MCE) and 40 ng/mL T3 for 3 days.

For the experiments about the differentiation efficiency effected by *Vim*, CG4 cells were seeded on 12-well plates at a density of 10^5^ cells per Matrigel-coated well and cultured for 1 day prior to transfection. The cells were then transfected with four different constructs, namely the Vector group, *Vim* KO+Vector group, *Vim* KO+*Vim*^WT^ group, and *Vim* KO+*Vim*^T35A^ ^T63A^ group. After transfection, the cells were subjected to bleomycin selection for 3 days, followed by an additional 3 days of differentiation induction. Then, immunostaining was performed to determine the proportion of MBP^+^ cells.

### Plasmid construction

Single guide RNAs (sgRNAs) were designed to target exon 1 of the mouse *Vim*, *Oga* and *Ogt* gene using the CRISPR design tool as described previously [40]. The selected sgRNA sequences for *Vim* (5’-GACACAGACCTGGTAGACA-3’ and 5’-CCCTTCGAAGCCATGTCTACC-3’), *Ogt* (5’-ACTGTCGGCCACGTTGCCCACGG-3’ and 5’-TAATAAGGCTGCAACCGGA-3’), and *Oga* (5’-TCAAGCGACGTTGGAGGAGCGGG-3’ and 5’-GCGAGAGATGTATTCAGTGGAGG-3’) were cloned into a pX330-gRNA vector that co-expressed Cas9 nuclease and an mCherry reporter. The full-length coding sequence of mouse *vimentin* (*Vim*, NM_011701.4) was amplified using polymerase chain reaction (PCR) from the total RNA extracted from mouse brains. The PCR products were then cloned into a pCDNA3.1 vector. To generate constructs for *Oga* and *Ogt* overexpression, the full-length coding sequences of mouse *Oga* (*Mgea5*, NM_023799) and *Ogt* (NM_139144) were amplified from mouse brain cDNA using PCR and cloned into the pCDNA3.1 vector.

To generate constructs for immunoprecipitation (IP) assays, we constructed the pCDNA3.1-Flag-*Vim*-HA plasmid. The full-length coding sequence of mouse *Vim* (NM_011701.4) was amplified with an N-terminal Flag tag and a C-terminal HA tag, and subsequently cloned into the pCDNA3.1 vector. Site-directed mutagenesis was performed on this template to generate the O-GlcNAc-deficient mutant pCDNA3.1-Flag-*Vim*^T35A^ ^T63A^-HA, in which Threonine 35 and Threonine 63 were substituted with Alanine.

### Cell transfection

Human embryonic kidney (HEK) 293T cells were maintained in DMEM supplemented with 10% FBS. For cell transfection, HEK293T cells were seeded at a density of 5×10^5^ cells per well in 6-well plates. At ∼70–80% confluence, cells were transfected with 3 mg plasmid DNA per well using polyethylenimine (PEI; Polysciences, Pennsylvania, USA). NPCs were transfected using the Neon Transfection System (Invitrogen, Waltham, USA), according to the manufacturer’s instructions.

### Protein sample preparation

For the O-GlcNAcome analysis, primary OPCs were isolated from a pool of 100 postnatal day 0 (P0) neonatal mice. Brains were rapidly dissected, and the brainstem, cerebellum, and midbrain were removed. The remaining tissue was digested using Trpzyme to create a single-cell suspension. OPCs were then purified from this suspension using the CD140α (Pdgfrα) MicroBead Kit (Miltenyi Biotec, Cologne, Germany) according to the manufacturer’s protocol. Total protein was extracted from OPCs using a lysis buffer containing 8 M urea and 1% protease inhibitor cocktail, sonicated three times on ice with a high-intensity ultrasonic processor (Scientz, Ningbo, China) in lysis buffer, and centrifuged at 12,000 g and 4°C for 10 min. The supernatant was collected, and protein concentration was determined using a BCA kit according to the manufacturer’s instructions.

### Liquid chromatography-tandem mass spectrometry (LC–MS/MS)

The O-GlcNAcome profiling was performed as a commercial service by PTM Bio (Hangzhou, China). To concentrate the proteins and remove interfering substances, acetone precipitation was performed. Briefly, one volume of pre-cooled acetone was added to the sample and vortexed, followed by the addition of four additional volumes of pre-cooled acetone. The mixture was incubated at -20°C for 2 h. The resulting protein precipitates were collected by centrifugation at 4,500 g for 5 min at 4°C. The supernatant was discarded, and the pellet was washed twice with pre-cooled acetone. Proteins were resuspended in digestion buffer and subjected to primary digestion with trypsin (1:50 trypsin-to-protein w/w ratio) overnight at 37°C. Subsequently, the sample was reduced with 5 mM dithiothreitol (DTT) for 60 min at 56°C and alkylated with 11 mM iodoacetamide (IAA) for 45 min at room temperature in the dark. A secondary digestion was then performed with trypsin (1:100 w/w) for 4 h at 37°C to ensure complete enzymatic cleavage.

Peptides were desalted using a C18 SPE column (activated with methanol and equilibrated with 0.1% TFA). Peptides were eluted with 80% acetonitrile (ACN) and quantified using a BCA assay. For the enrichment of O-GlcNAc-modified peptides, desalted peptides were redissolved in NETN buffer (100 mM NaCl, 1 mM EDTA, 50 mM Tris-HCl, 0.5% NP-40, pH 8.0) and incubated with O-GlcNAc antibody-conjugated beads (PTM-954, PTM Bio) at 4°C overnight with gentle rotation. The immunocomplexes were washed four times with NETN buffer and twice with double-distilled water. The enriched glycopeptides were eluted and subjected to a final cleanup using C18 ZipTips to remove trace salts prior to LC-MS/MS analysis.

The tryptic peptides were dissolved in solvent A (0.1% formic acid, 2% acetonitrile in water), directly loaded onto a custom-made reversed-phase analytical column (25-cm length, 75/100 μm i.d.). Peptides were separated with a gradient from 6% to 24% solvent B (0.1% formic acid in acetonitrile) over 70 min, 24% to 35% in 14 min and climbing to 80% in 3 min then holding at 80% for the last 3 min, all at a constant flow rate of 450 nL/min on a nanoElute UHPLC system (Bruker Daltonics). The peptides were subjected to capillary source followed by the timsTOF Pro (Bruker Daltonics) mass spectrometry. The electrospray voltage was set to 1.60 kV. Precursors and fragments were analyzed at the TOF detector, with a MS/MS scan range from 100 to 1700 m/z. The timsTOF Pro was operated in parallel accumulation serial fragmentation (PASEF) mode. Precursors with charge states 0 to 5 were selected for fragmentation, and 10 PASEF-MS/MS scans were acquired per cycle. The dynamic exclusion was set to 30 s.

The raw MS/MS data were processed using the MaxQuant search engine (v1.6.15.0). Tandem mass spectra were searched against the Mus_musculus_10090_SP_20210721.fasta database concatenated with a reverse decoy database. Trypsin/P was specified as the cleavage enzyme allowing up to 2 missing cleavages. The mass tolerance for precursor ions was set to 20 ppm for both first and main searches, and the fragment ion mass tolerance was 20 ppm. Carbamidomethylation on Cys was set as a fixed modification. Acetylation (protein N-terminal), oxidation (Met), and O-glycosylation (+203.08 Da on Ser/Thr) were specified as variable modifications. False discovery rate (FDR) of protein, peptide and PSM was adjusted to < 1%.

### Western blotting

Cells were lysed in RIPA buffer (Beyotime, Xiamen, China) supplemented with phenylmethanesulfonyl fluoride (PMSF; Beyotime) and Thiamet-G (APExBIO, Houston, USA), homogenized for 30 min on ice and centrifuged at 12000 rpm for 30 min. Protein samples (10 mg) were separated using sodium dodecyl sulfate-polyacrylamide gel electrophoresis (SDS-PAGE) gels and transferred to polyvinylidene fluoride (PVDF; Millipore, Massachusetts, USA) membranes. Membranes were blocked in 5% skim milk at room temperature for 1 hour and then incubated with primary antibody for O-GlcNAc (Invitrogen), β-Actin (Abclonal, Wuhan, China) (1:1000 dilution) at 4°C overnight. Membranes were consequently washed using TBST three times and then incubated with the appropriate HRP-conjugated secondary antibodies (Abclonal) at room temperature for 1 hour. Bands were visualized using a luminescent image analyser (GE Imagination LAS 4000; GE imagination at work, Boston, USA). The results of protein band intensity were quantified with ImageJ (NIH, Bethesda, USA) software.

### Immunoprecipitation

Cells expressing Flag-HA-tagged *Vim*^WT^ and *Vim*^T35A^ ^T63A^ were lysed in IP lysis buffer containing protease inhibitors and Thiamet-G. Lysates were incubated with anti-Flag magnetic beads (Bimake, Texas, USA) overnight at 4°C with rotation. The beads were washed extensively with lysis buffer, and the bound protein complexes were eluted by boiling in SDS loading buffer. The eluates were subsequently analyzed by Western blotting using anti-O-GlcNAc (1:1000; PTM bio), anti-Flag (1:10000; Sigma-Aldrich, Melbourne, Australia) and anti-HA (1:10000; Abclonal) antibodies to evaluate the O-GlcNAcylation levels of Vimentin.

### Quantitative real-time PCR (qRT-PCR)

Total RNA was extracted from cultured NPCs, OPCs, and OLs at various stages of differentiation using TRNzol Universal Reagent (TIANGEN, Beijing, China) according to the manufacturer’s protocol. Briefly, cells were lysed directly in the culture dish, and RNA was isolated following extraction using chloroform and precipitation using isopropanol. The RNA pellet was washed with 75% ethanol, air-dried, and resuspended in RNase-free water. The quantity and purity of the RNA were assessed using a NanoDrop 2000 spectrophotometer (Thermo Fisher Scientific, Waltham, USA); samples having an A260/280 ratio between 1.8 and 2.0 were used for downstream applications.

First-strand cDNA was synthesized from 1 µg DNase-treated total RNA using a HiScript II1st Strand cDNA Synthesis Kit (Vazyme, Nanjing, China), which included a blend of oligo(dT) and random hexamer primers to ensure unbiased representation of the transcriptome.

qRT-PCR was performed on a CFX96 Real-Time PCR Detection System (Bio-Rad, Hercules, USA) using ChamQ SYBR Color qRT-PCR Master Mix (Vazyme). Each 10 µL reaction mixture contained 5 µL 2X ChamQ SYBR qRT-PCR Master Mix, 200 nM each forward and reverse primer, and 1 µL diluted cDNA template. Thermal cycling conditions were as follows: initial denaturation at 95°C for 3 minutes, followed by 40 cycles at 95°C for 15 seconds and at 60°C for 60 seconds. A melt curve analysis was performed at the end of each run to verify the specificity of the amplification product. Gene expression levels were quantified using comparative C[(2^−ΔΔCt^) method with expression normalized to that of the housekeeping gene β-actin (*Actb*). Primers used for qRT-PCR: *Vim* (F: CGTCCACACGCACCTACAG; R: GGGGGATGAGGAATAGAGGCT), *Actb* (F: CTGAATCCATCCCAGTAGCCT; R: CTGAATCCATCCCAGTAGCCT); *Dbn1* (F: GCTTGCAGCGTCAGGAGAA; R: TCCTCACCAACCCAGTTGATG); *Flnb* (F: GACCACGAAAGCATTAAGCTCG; R: CTGAATCCATCCCAGTAGCCT); mature OL markers including *Mbp* (F: TCCATCCCAAGGAAAGGGGA; R: TCTGCCTCCGTAGCCAAATC), *Plp1* (F: CTGCAAAACAGCCGAGTTCC; R: ATCAGAACTTGGTGCCTCGG), *Mog* (F: CCTTCAGGCTTCTTGGGGTA; R: TCACTCTGAACTGTCCTGCG), and *Mag* (F: GCCAGACCATCCAACCTTCT; R: CCTCACCTCAGTCTCCCTGA); OPC markers including *Pdgfr*α (F: AGAGTTACACGTTTGAGCTGTC; R: GTCCCTCCACGGTACTCCT) and *Olig2* (F: GGGAGGTCATGCCTTACGC; R: CTCCAGCGAGTTGGTGAGC).

### Immunostaining and imaging

OLs and HEK293T cells were fixed with 4% PFA in PBS for 10 minutes at room temperature, washed three times with PBS, permeabilized with 0.2% Triton X-100 in PBS for 10 minutes, and blocked with 7.5% BSA in PBS for 1 hour. Cells were incubated overnight at 4°C with primary antibodies against O-GlcNAc (1:500; Abcam, Cambridge, UK), O4 (1:500; R&D Systems), Pdgfrα (1:500; Oasis, Hangzhou, China), or HA (1:500; Abclonal) in blocking buffer, followed by Alexa Fluor-conjugated secondary antibodies (1:1000; Invitrogen) and DAPI for 1 hour at room temperature. Tissue sections were blocked with 5% donkey serum in PBS with 0.1% Triton X-100 for 1 hour, incubated with primary antibodies against O-GlcNAc (1:500; Abcam), CC1 (1:500; Oasis), or Pdgfrα (1:500; Oasis) overnight at 4°C, and then with secondary antibodies for 2 hours. Imaging was performed using a Nikon ECLIPSE Ti2 (Nikon, Melville, USA).

### Cell proliferation and apoptosis assays

To assess cell proliferation, transfected CG4 cells were fixed with 4% PFA and subjected to immunofluorescence staining. Cells were incubated with primary antibodies against Ki67 (1:500; MCE) and Sox10 (1:500; Oasis) overnight at 4°C, followed by incubation with corresponding secondary antibodies. The proliferation rate was quantified as the percentage of Ki67[cells within the Sox10[cell population.

For the assessment of apoptosis, the Terminal Deoxynucleotidyl Transferase dUTP Nick End Labeling (TUNEL) assay was performed using a TUNEL BrightGreen Apoptosis Detection Kit (Vazyme) according to the manufacturer’s instructions. Cells were simultaneously co-stained with an anti-Pdgfrα antibody (1:500; Oasis) to identify OPCs. The apoptotic rate was calculated as the ratio of TUNEL[Pdgfrα[cells to the total Pdgfrα[population. Images were acquired and analyzed using the same microscopy and software settings as described above.

### Image quantification

All fluorescence images were analyzed using ImageJ. To ensure accurate interpretation of O-GlcNAc signal in OPCs and oligodendrocytes, we measured mean fluorescence intensity (MFI) exclusively in cells positively identified by lineage-specific markers (Pdgfrα for OPCs; O4 for OLs; CC1 for more mature oligodendrocytes; and Mbp for functional oligodendrocytes). This per-cell analysis eliminates the need for physical isolation and avoids potential confounding signals from non-lineage cells. Specifically, individual marker-positive cells were segmented based on their immunostaining, and O-GlcNAc MFI was quantified strictly within these cellular boundaries.

### Proteomics data analysis and functional annotation

Gene Ontology (GO) and Kyoto Encyclopedia of Genes and Genomes (KEGG) pathway analyses were performed using DAVID [41]. Bubble plots were generated using an online platform (https://www.bioinformatics.com.cn, accessed December 10, 2024) [42]. Protein localization was predicted using WoLF PSORT [43].

For the analysis of O-GlcNAcylation motifs, 21-amino-acid sequences (±10 residues) flanking identified sites were analyzed using OmicShare tools [44]. Intrinsically disordered regions and linear interacting peptides were identified using MobiDB [45], and protein domains were annotated using the Pfam database [46].

### Protein-protein interaction (PPI) network analysis

The list of O-GlcNAcylated proteins was submitted to STRING (https://string-db.org) to extract known and predicted functional associations (confidence score of > 0.4) and visualize the results using Cytoscape (V3.10.1).

### Molecular docking simulations

Molecular docking was performed using HPEPDOCK [47] with the human OGT crystal structure (PDB ID: 4GYW) as the receptor [48, 49]. Structures were prepared in PyMOL (v3.1.1) by removing water molecules, adding polar hydrogens, and assigning Gasteiger charges. The interface areas and ΔiG between OGT and the peptides were calculated using PDBePISA (https://www.ebi.ac.uk/pdbe/pisa/).

### Statistical analysis

Error bars represent mean ± SD from at least three independent experiments. Statistical analyses were performed using GraphPad Prism 9 (GraphPad Software, San Diego, USA). We explicitly define that for all assays “n” represents the number of independent biological replicates. In brief, for *in vitro* cell culture assays, “n” indicates the number of independent experiments, each with three technical replicates; for *in vivo* experiments, “n” denotes the number of mice per group, with samples collected from individual mice processed as separate biological replicates. Comparisons between two groups were performed using a two-tailed unpaired Student’s t-test. Multiple-group comparisons were performed using one-way ANOVA with Tukey’s post-hoc test as appropriate. Specific statistical tests used are clearly stated in each figure legend. No statistical methods were used to predetermine sample size.

## Results

### Global O-GlcNAcylation levels in OPCs

To investigate the role of O-GlcNAcylation in OPC lineage progression, we first characterized global O-GlcNAc levels during the transition from OPCs to mature OLs. To this end, primary neural precursor cells (NPCs) isolated from embryonic day 12.5 (E12.5) mouse cortices were cultured under established differentiation conditions (Supplementary Figure S1A, B). Immunofluorescence showed approximately 20% Pdgfrα^+^ OPCs at the precursor stage, and 12% of Mbp^+^ OLs following differentiation (Supplementary Figure S1C, D). Both immunostaining and Western blot analyses revealed an upregulation of global O-GlcNAcylation in OLs relative to OPCs. (Figure 1A-D). Quantitative analysis of *in vitro* cultures showed that the O-GlcNAc mean fluorescence intensity (MFI) higher in O4^+^ OLs (45.4) than in Pdgfrα^+^ OPCs (5.2) (Figure 1A, B). This protein-level increase was further confirmed by Western blotting, which exhibited a 1.3-fold increase in global O-GlcNAcylation during differentiation (Figure 1C, D). Consistent with these changes, qRT-PCR analysis demonstrated a 1.2-fold induction of *Ogt* mRNA as cells transitioned from the precursor stage to mature OLs (Figure 1E).

**Figure 1.**
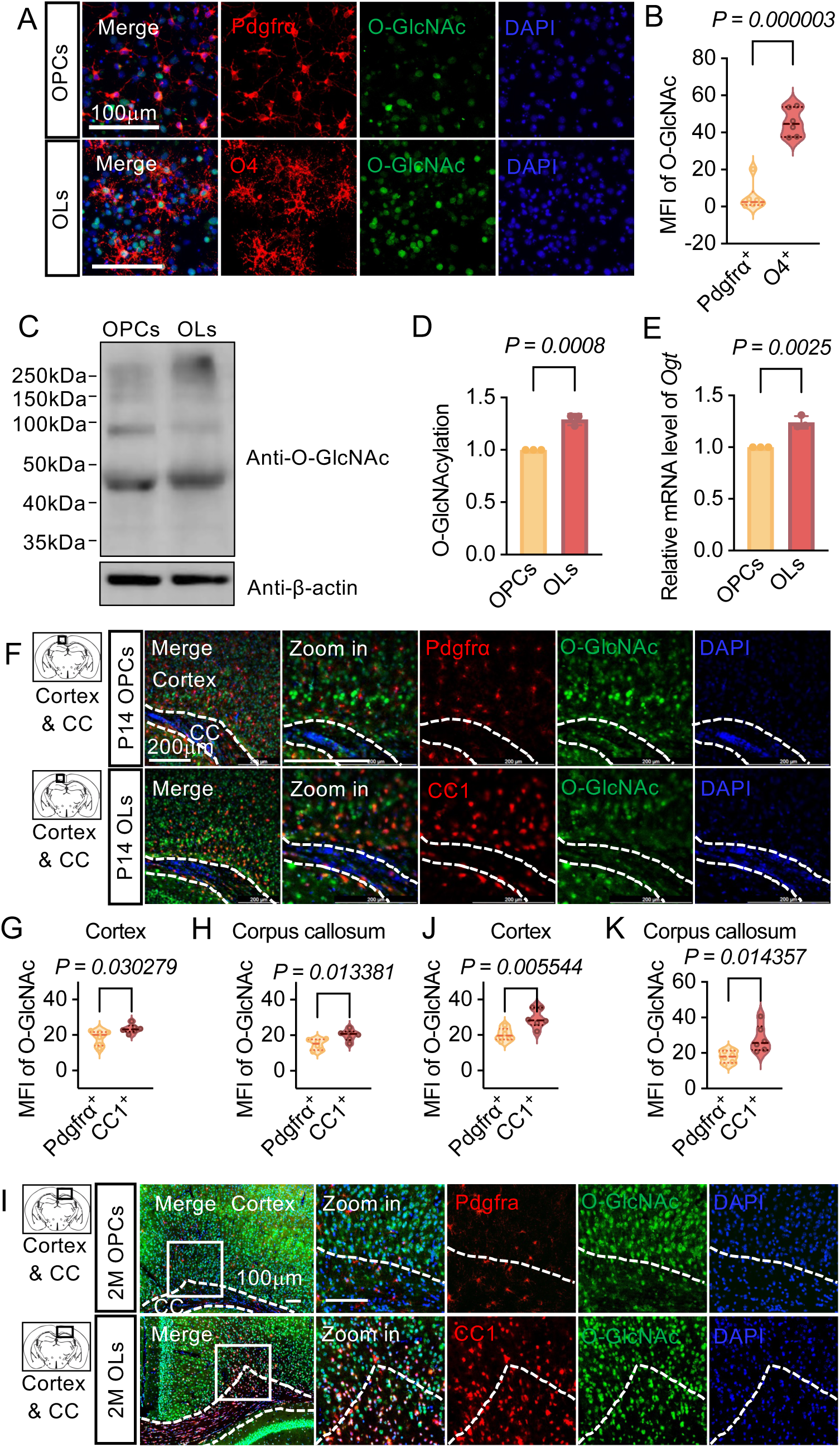
O-GlcNAcylation levels in the oligodendrocyte lineage. (A) Representative immunostaining images of oligodendrocyte precursor cells (OPCs; Pdgfrα[, red) and differentiating oligodendrocytes (OLs; O4[, red) differentiated from primary neural precursor cells (NPCs). Staining shows O-GlcNAcylation (O-GlcNAc^+^, green) and nuclei (DAPI, blue). Scale bar: 100 μm. n=6 independent biological experiments. (B) Quantification of the mean fluorescence intensity (MFI) of O-GlcNAc in Pdgfrα[OPCs and O4[OLs from (A). (C, D) Western blot (C) and quantification (D) of total O-GlcNAcylation levels in OPCs and OLs. n=3 independent biological experiments. (E) qRT-PCR analysis of O-GlcNAc transferase (*Ogt*) mRNA expression in OPCs and OLs. (F) *In vivo* immunolabeling of OPCs (Pdgfrα[, red) and mature OLs (CC1[, red) in the brains of P14 mice, showing O-GlcNAcylation levels (O-GlcNAc^+^, green). Scale bar: 200 μm. n=6 mice per group. (G, H) Quantification of the MFI of O-GlcNAc in *in vivo* Pdgfrα[OPCs and CC1[OLs in cortex (G) and corpus callosum (CC) (H). (I-K) *In vivo* immunostaining images (I) and quantification (J, K) of O-GlcNAcylation levels in Pdgfrα[OPCs and CC1[OLs in the 2 months old (2M) mouse brain. Scale bar: 100 μm. n=6 mice per group. All quantifications were performed on a per-cell basis. *P*-values were calculated using two-tailed unpaired Student’s t-test.

To validate these findings *in vivo*, we performed immunostaining on brain sections from postnatal day 14 (P14) and 2-month-old (2M) adult mice, focusing on the corpus callosum (CC) and cortex (Figure 1F, I). Consistent with our *in vitro* data, O-GlcNAc levels (MFI) at P14 were lower in Pdgfrα^+^ OPCs (18.6) than in CC1^+^ OLs (23.6) within the cortex, while in the CC, the intensity increased from 14.8 in OPCs to 20.1 in OLs (Figure 1G, H). A similar trend was observed in the 2M adult brain, with MFI rising from 20.4 in OPCs to 29.4 in OLs in the cortex, and from 17.9 to 28.0 in the CC (Figure 1J, K). Density plots of MFI revealed a consistent shift in the distribution curve for OLs compared to OPCs across both ages and regions (Figure 1G, H, J, K).

To determine the functional requirement for O-GlcNAc cycling in lineage progression, we genetically modulated the expression of O-GlcNAc cycling enzymes in primary E12.5 NPCs. Primary NPCs were transfected with plasmids for either the overexpression or CRISPR/Cas9-mediated knockout of *Ogt* or *Oga*, followed by Bleomycin selection and differentiation. Immunostaining analysis revealed that decreasing global O-GlcNAcylation, achieved through either *Oga* overexpression or *Ogt* knockout, impaired the transition from OPCs to OLs (Supplementary Figure S2A, D). Specifically, *Oga* overexpression reduced the proportion of O4^+^ OLs from approximately 7% in control cultures to around 2%, while Mbp^+^ cells decreased from approximately 4% to 1% (Supplementary Figure S2B, C). Mirroring these results, *Ogt* knockout similarly impaired the transition, with O4^+^ and Mbp^+^ percentages dropping to 1.8% and 0.5%, respectively (Supplementary Figure S2E, F). Conversely, augmenting O-GlcNAc levels via *Ogt* overexpression or *Oga* knockout facilitated the differentiation process. Overexpression of *Ogt* promoted maturation, increasing the percentage of O4^+^ cells to approximately 10% and Mbp^+^ cells to approximately 6% compared to empty vector controls (Supplementary Figure S2B, C). Furthermore, *Oga* knockout led to an increase in O4^+^ and Mbp^+^ populations to 7% and 5.5%, respectively (Supplementary Figure S2E, F). Collectively, these results demonstrate that O-GlcNAcylation is critical for OPC differentiation.

### Proteomic profiling of O-GlcNAcome in OPCs

To comprehensively characterize the O-GlcNAc landscape in OPCs, we conducted a proteomic analysis. Primary OPCs were isolated from postnatal day 0 (P0) mouse brains, followed by the enrichment of O-GlcNAcylated proteins using a combinatorial approach involving O-GlcNAc-specific antibody-conjugated beads and metal affinity chromatography (Figure 2A). The high purity of the isolated primary OPCs, determined to be 91.8% via flow cytometry, ensured the reliability and fidelity of the downstream proteomic dataset (Supplementary Figure S3). Enriched O-GlcNAcylated peptides were analyzed using Liquid Chromatography-Tandem Mass Spectrometry (LC-MS/MS). This analysis processed 15,932 total spectra, yielding 3,683 matched spectra and 3,065 peptides, which ultimately identified 165 O-GlcNAcylation sites mapping to 118 distinct proteins (Figure 2B). Subcellular localization analysis revealed that the OPC O-GlcNAcome was predominantly nuclear (64.4%), with substantial representation in the cytoplasmic (15.3%) and mitochondrial (4.3%) (Figure 2C). Notably, 78.8% of the identified proteins harbored a single O-GlcNAcylation site, while 21.2% contained two or more sites (Figure 2D). We cross-referenced our dataset with the O-GlcNAc Database and O-GlcNAcAtlas (v3.0) [38, 39]. This comparative analysis identified 74 novel O-GlcNAcylation sites (44.8%) on 67 proteins (56.8%) within our OPC dataset. Strikingly, 22 of these proteins had not been previously reported to undergo O-GlcNAcylation in any biological context (Figure 2E, F).

**Figure 2.**
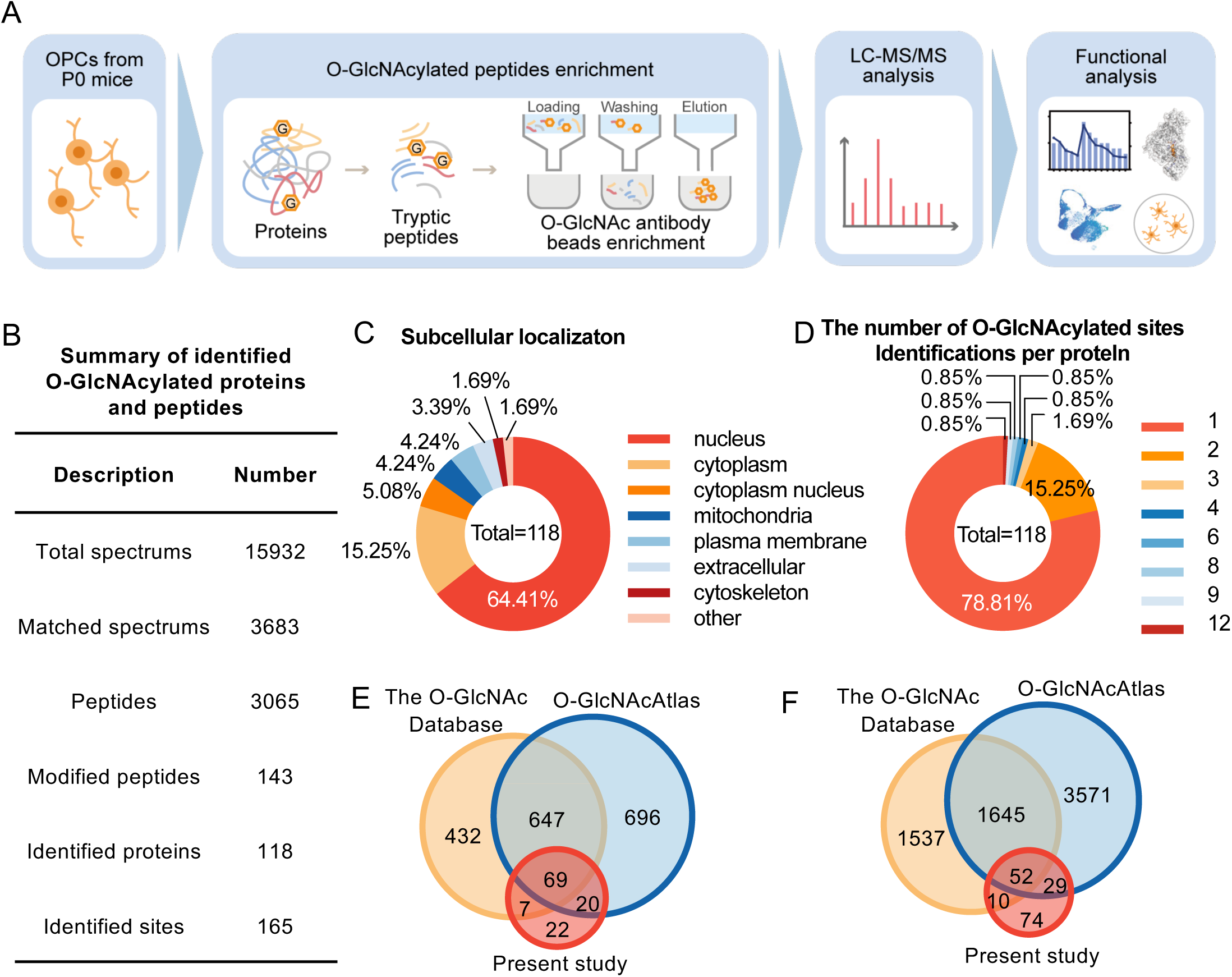
Proteomic analysis of O-GlcNAcylated proteins in OPCs. (A) Schematic of the experimental workflow for identifying O-GlcNAcylated proteins in OPCs, including OPC isolation from P0 mouse brains, protein extraction, O-GlcNAcylated peptides enrichment, LC-MS/MS analysis, and functional analysis. (B) Summary of the proteomics results, detailing the number of total spectra, matched spectra, peptides, modified peptides, identified proteins, and sites. (C) Pie chart showing the subcellular localization of the 118 identified O-GlcNAcylated proteins. (D) Distribution of O-GlcNAcylated sites per protein. (E, F) Venn diagrams comparing the identified O-GlcNAcylated proteins (E) and sites (F) from this study with the O-GlcNAc Database and O-GlcNAcAtlas, revealing 22 novel O-GlcNAcylated proteins and 74 novel modification sites.

### Functional annotation of the OPC O-GlcNAcome

We performed a functional enrichment analysis of O-GlcNAcylated proteins in OPCs. Gene Ontology (GO) analysis revealed a significant enrichment of these proteins in biological processes essential for OPC lineage progression (Figure 3A). Cellular component analysis indicated that the O-GlcNAcome is primarily localized to the cell body and cytoplasm (Figure 3B). Molecular function categories were enriched in protein binding and protein domain-specific binding (Figure 3C). Kyoto Encyclopedia of Genes and Genomes (KEGG) pathway analysis further revealed that these O-GlcNAcylated proteins are involved in gap junction signaling. Additionally, several neurodegenerative pathways were enriched, including those associated with Alzheimer’s, Parkinson’s, and Huntington’s diseases (Figure 3D). To investigate the topological organization of these modified proteins, we constructed a protein-protein interaction (PPI) network. This analysis identified several nodes, including Actb, Mapt, Tubb3, Htt, Ywhaq, and Srcin1 (Figure 3E). These proteins were reported to coordinate cytoskeletal dynamics and intracellular transport, both of which are essential processes underpinning the migration and differentiation of OPCs [9, 50–54]. Collectively, these findings provide a functional framework for the OPC O-GlcNAcome.

**Figure 3.**
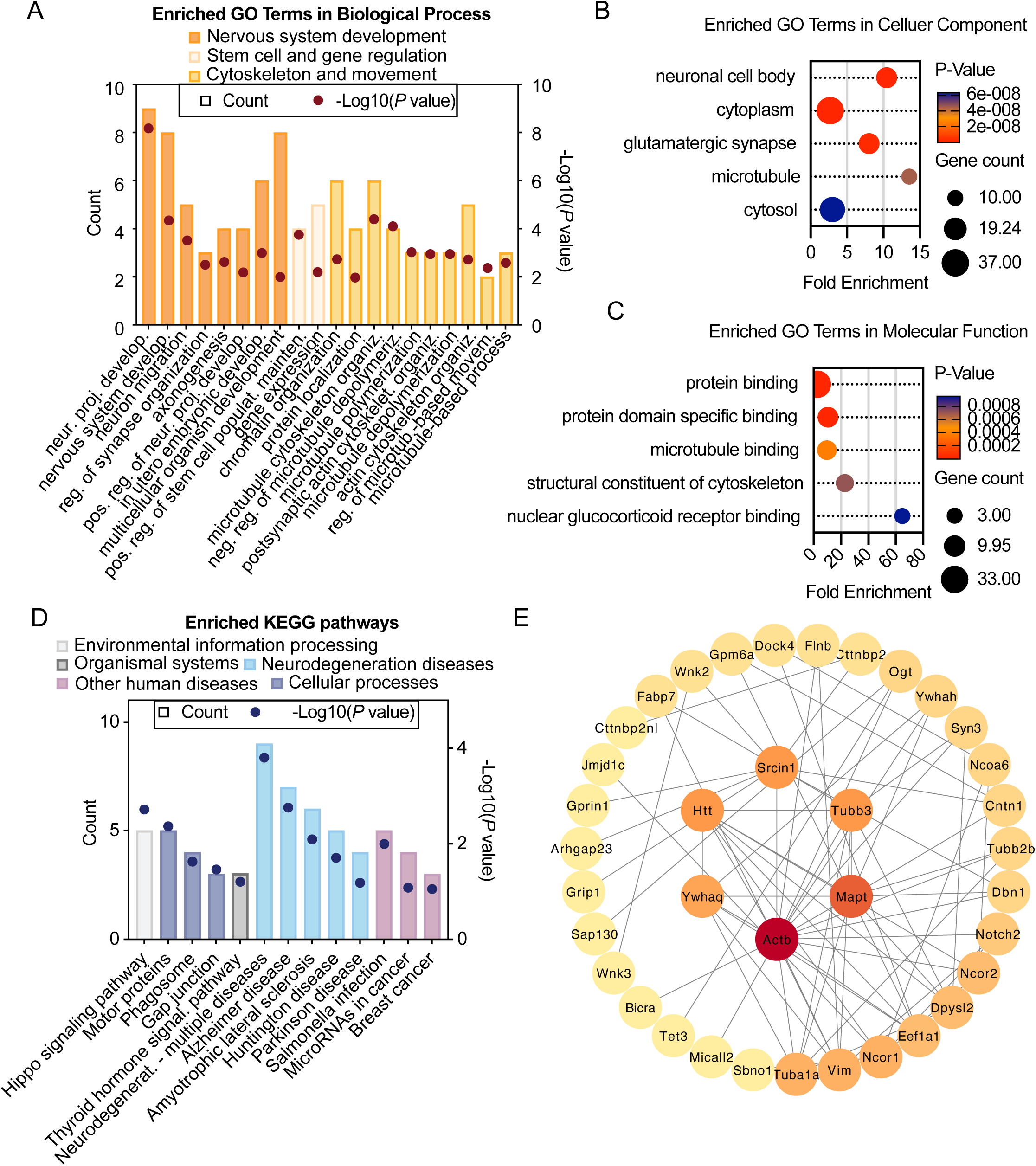
Functional annotation of the OPC O-GlcNAcylated proteins. (A) Gene Ontology (GO) enrichment analysis of the OPC O-GlcNAcylated proteins. (B, C) Bubble chart of enriched GO terms in cellular component (B) and molecular function (C) for O-GlcNAcylated proteins. (D) KEGG pathway analysis reveals associations with pathways linked to neurodegenerative diseases including Alzheimer’s disease. (E) Protein-protein interaction network of the OPC-specific O-GlcNAcylated proteins.

### Motif and structural analysis of OPC O-GlcNAcylated proteins

To characterize the sequence and structural determinants underlying O-GlcNAcylation in OPCs, we conducted a motif analysis on the 165 identified O-GlcNAcylation sites. This analysis revealed a distinct preference for proline at the -3 and -2 position, valine at the -1 position relative to the modified residue, arginine at the +1 position, with serine and alanine enriched in the flanking regions (Figure 4A). These features are consistent with the known structural roles of proline in imparting conformational rigidity and stabilizing binding pockets, while distal arginine residues may facilitate downstream protein-protein interactions [55, 56]. Previous studies have established that Ogt modification sites are predominantly localized within IDRs [34, 57], which is essential for dynamic protein-protein interactions, molecular recognition, and cellular signaling regulation [58]. We further integrated structural predictions from MobiDB, covering intrinsically disordered regions (IDRs) and linear interacting peptides (LIPs), and Pfam for structured domains to contextualize these modification sites. Of the all 165 O-GlcNAcylation sites identified, the majority (69.7%, n=115) were localized within IDRs, while 35.2% (n=58) were situated within LIPs. In contrast, only a minority (19.4%, n=32) of sites mapped to ordered domains (Figure 4B-D). Functional enrichment analysis revealed distinct biological roles based on structural context. Proteins modified within IDRs showed strong enrichment in synapse organization and regulation of protein-containing complex assembly (Figure 4B). Proteins modified at LIP sites were linked to synapse organization, negative regulation of vascular endothelial growth factor signaling pathway and epidermal growth factor receptor signaling pathway (Figure 4C). The limited number of sites within structured domains were associated with cytoskeletal regulation (Figure 4D).

**Figure 4.**
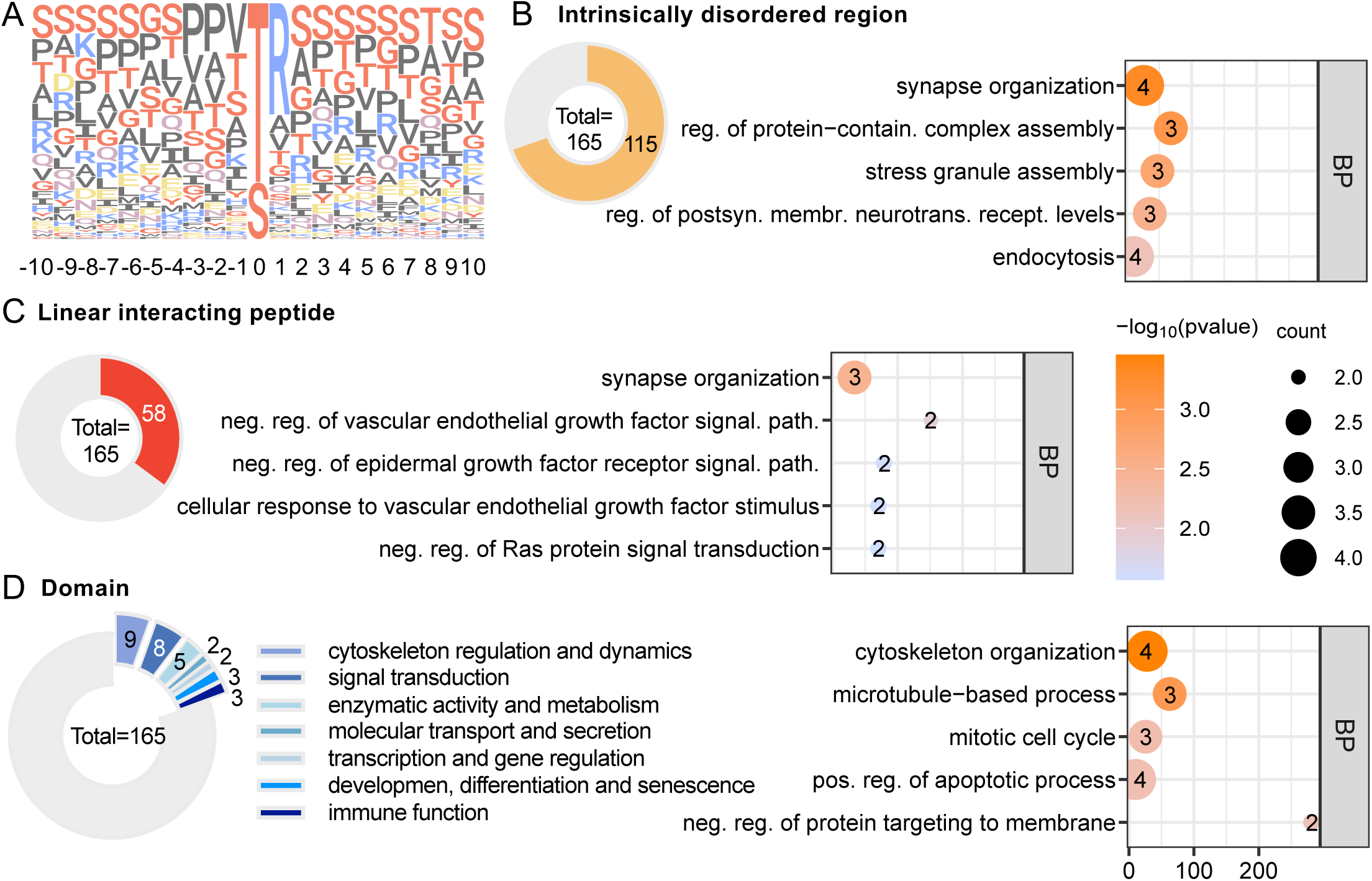
Motif and structural analysis of OPC O-GlcNAcylated sites. (A) Motif analysis of the amino acid sequences surrounding the OPC O-GlcNAcylated sites. (B) Distribution of O-GlcNAcylated sites within intrinsically disordered regions (IDRs) and GO enrichment analysis of the corresponding proteins. (C) Distribution of O-GlcNAcylated sites within linear interaction peptides (LIPs) and GO enrichment analysis of the corresponding proteins. (D) Distribution of O-GlcNAcylated sites within structured protein domains and the functional classification of the corresponding proteins.

To highlight the unique regulatory profile of the oligodendrocyte lineage, we performed a parallel analysis focusing exclusively on the 74 newly identified O-GlcNAcylation sites (Supplementary Figure S4A, C-E). Remarkably, the structural distribution of these sites showed a similar preference, with a predominant localization in IDRs (64.9%) and LIPs (35.1%). However, these OPC-specific modifications were more specifically enriched in pathways essential for CNS development and cytoskeletal organization.

### Chemical properties of O-GlcNAcylation motifs in OPCs

To characterize the chemical microenvironment governing Ogt substrate recognition, we analyzed the amino acid residues flanking the 165 identified O-GlcNAcylation sites (Figure 5A). Small, nonpolar, and hydrophobic residues prevailed in the immediate vicinity, specifically from positions -3 to +3, while basic residues were enriched at the +1 position. Notably, the 74 novel O-GlcNAcylation sites exhibited identical chemical preferences to the global profile (Supplementary Figure S4B). This prevalence of hydrophobic residues suggests that O-GlcNAcylation frequently occurs at protein-protein interfaces, potentially modulating interaction specificity and complex stability [59].

**Figure 5.**
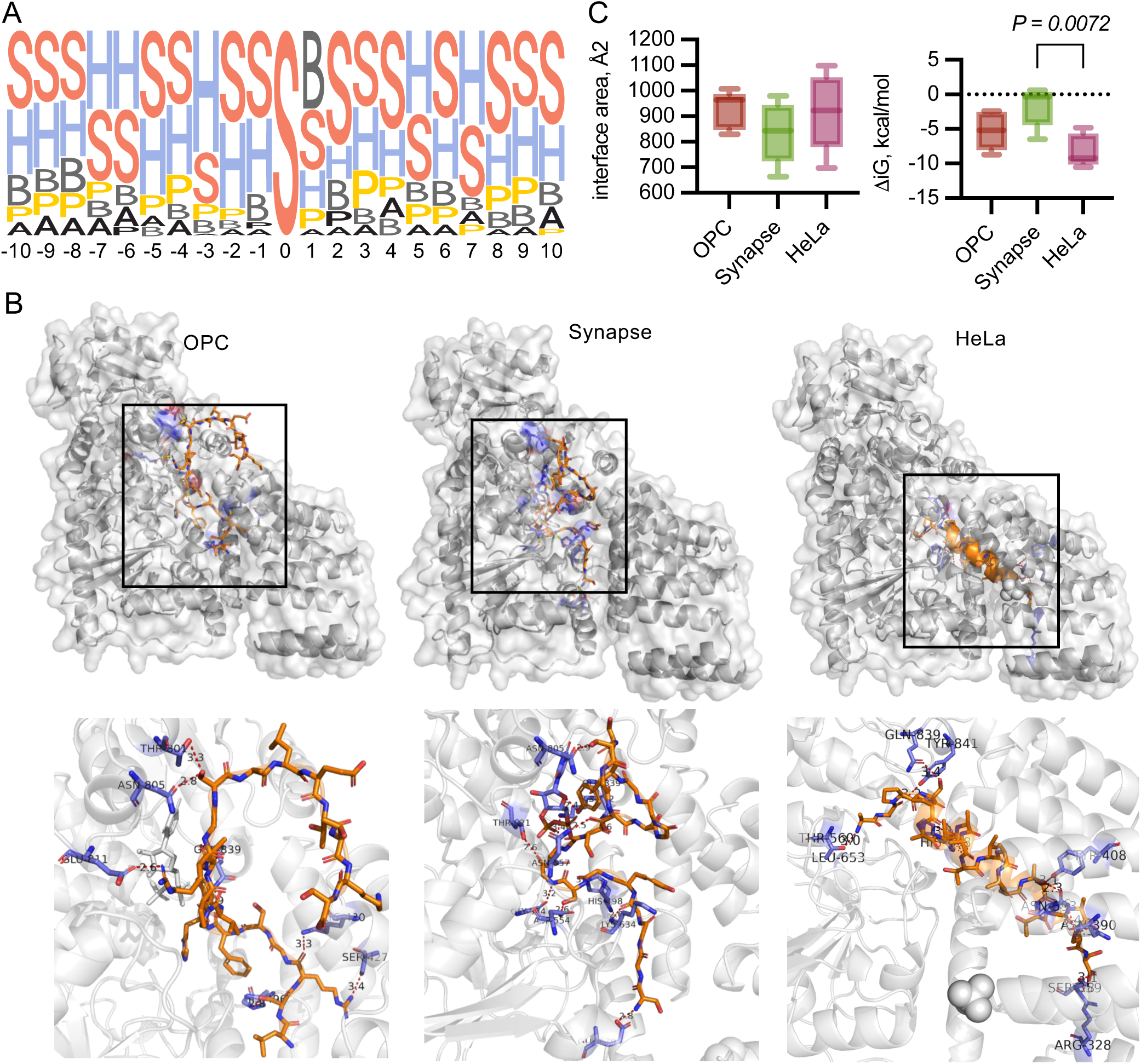
Chemical properties and OGT Interactions of OPC O-GlcNAcylation motifs. (A) Analysis of the chemical properties of amino acids surrounding O-GlcNAcylated sites in OPCs. (B) Molecular docking simulations showing the binding modes of representative O-GlcNAcylation peptides derived from OPCs, synapses, and HeLa cells with O-GlcNAc transferase (OGT). (C) Comparison of typical motifs, interface area, and binding free energy (ΔG) for OPCs, synapses, and HeLa cells from (B). *P*-values were calculated using one-way ANOVA test.

Furthermore, the presence of specific polar and ionizable residues likely facilitates the stabilization of the Ogt-substrate complex through polar-polar contacts within these hydrophobic pockets [56]. To evaluate the energetic landscape of Ogt binding, we performed molecular docking simulations using representative O-GlcNAcylated peptides from OPCs, synapses, and HeLa cells [34, 35]. These simulations revealed that OPC-derived peptides exhibit intermediate binding free energies (ΔG) compared to their synaptic and HeLa-derived counterparts (Figure 5B, C). Such intermediate energetics may allow for reversible and dynamic modulation of the O-GlcNAcome in response to fluctuating metabolic cues throughout the oligodendrocyte lineage.

### Association of OPC O-GlcNAcylated proteins with multiple sclerosis

To further investigate the potential involvement of O-GlcNAcylated proteins in demyelinating pathologies, we intersected these O-GlcNAc-modified proteins with two multiple sclerosis-related resources: a single-nucleus RNA sequencing (snRNA-seq) dataset, from which we computationally isolated the OPC cluster, and a proteomic dataset derived from multiple sclerosis lesions [60, 61]. This integrative analysis identified 15 O-GlcNAcylated proteins shared across all datasets (Supplementary Figure S5A; Table 1). These genes are enriched in biological processes related to cytoskeleton organization (Supplementary Figure S5B). Subcellularly, these proteins were predominantly localized to the cytoplasm and cell projection (Supplementary Figure S5C). Molecular function analysis further indicated a strong enrichment in profilin binding, which underscores the role of these targets in facilitating molecular interactions essential for OPC function (Supplementary Figure S5D). Furthermore, KEGG pathway analysis revealed that these O-GlcNAcylated candidates are significantly enriched in four functional pathways: IgSF CAM signaling, microRNAs in cancer, Alzheimer’s disease, and pathways of neurodegeneration (Supplementary Figure S5E). Among the overlapping candidates, Vimentin (Vim), Drebrin 1 (Dbn1), and Filamin B (Flnb) exhibited the most pronounced expression changes (|log_2_FC| > 1.5 in at least one dataset) in people with multiple sclerosis (Supplementary Figure S5F). Therefore, we focused our study on these candidates.

**Table 1.** OPC O-GlcNAcylated genes shared with both multiple sclerosis lesions proteome and multiple sclerosis snRNA-seq datasets.

| **Gene name** | **Subcellular localization** | **Number of O-GlcNAcylated sites** | **Number of OPC-specific O-GlcNAcylated sites** |
| --- | --- | --- | --- |
| *Apc* | cytoplasm | 1 | 1 |
| *Dbn1* | cytoplasm | 2 | 2 |
| *Dmtn* | plasma membrane | 1 | 0 |
| *Flnb* | cytoplasm | 1 | 1 |
| *Htt* | cytoplasm | 1 | 1 |
| *Irs2* | cytoplasm | 1 | 0 |
| *Kif26b* | cytoplasm | 1 | 1 |
| *Map6* | cytoplasm | 1 | 0 |
| *Mtap* | cytoplasm | 1 | 1 |
| *Mtss2* | cytoplasm | 1 | 1 |
| *Pclo* | cell projection | 6 | 0 |
| *Sgip1* | membrane | 1 | 0 |
| *Trio* | cytoplasm | 1 | 0 |
| *Tubb2b* | cytoplasm | 1 | 1 |
| *Vim* | cytoplasm | 2 | 2 |

### Expression dynamics of OPC O-GlcNAcylated proteins during aging

To investigate the potential role of O-GlcNAcylation in the context of aging, we first measured total O-GlcNAc levels in Pdgfrα^+^ OPCs in mouse brains at P14, 2 months, and 12 months (Supplementary Figure S6A). We found that total O-GlcNAc levels in OPCs gradually increased from postnatal development to young adulthood and subsequently declined with advanced aging in both the cortex and the CC. Specifically, the MFI rose from 22.1 at P14 to 29.4 at 2M in the cortex and from 19.3 to 28.0 in the CC, followed by a decrease to 18.8 and 13.0, respectively, at 12M (Supplementary Figure S6B, C).

To explore whether key O-GlcNAcylated candidates contribute to age-related deficits in myelin repair capacity, we analyzed the expression dynamics of *Vim*, *Dbn1*, and *Flnb* in OPCs across the mouse lifespan. Transcriptomic data revealed that the expression of *Vim*, *Dbn1*, and *Flnb* peaked during early development, specifically from P0 to P7, and declined thereafter (Figure 6A). In particular, *Vim* showed robust expression in neonatal OPCs followed by a decline during aging. *Dbn1* expression reached its maximum at P7 before decreasing, while *Flnb* exhibited high expression at P0 but remained low throughout later life stages.

**Figure 6.**
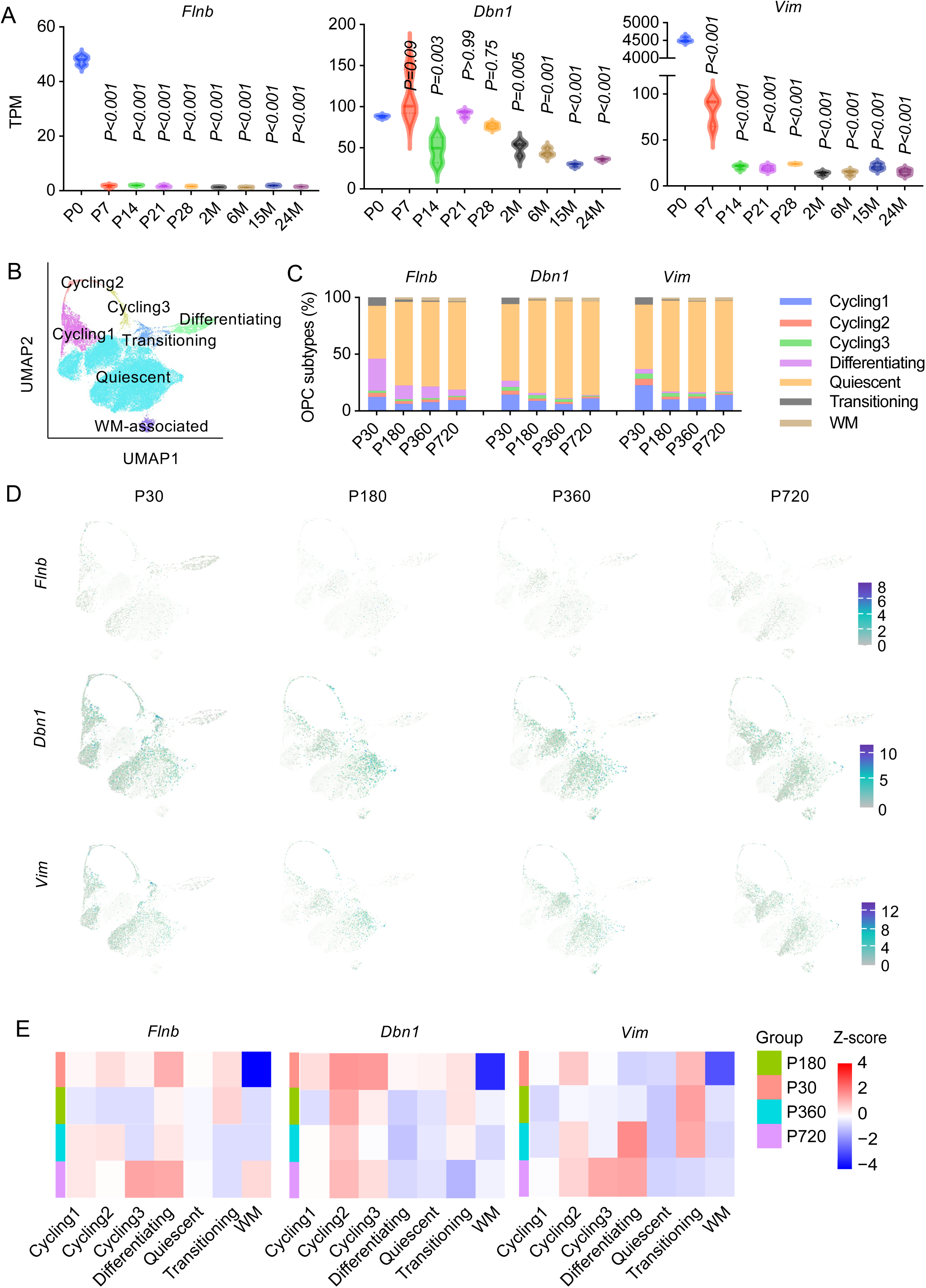
Expression dynamics of *Vim*, *Dbn1*, and *Flnb* in OPC subtypes during aging. (A) Transcriptomic data (TPM) showing significant expression changes of *Flnb*, *Dbn1*, and *Vim* across postnatal ages (P0 to 24M). (B) UMAP plot illustrating OPC subtypes (cycling, differentiating, quiescent, and transitioning) identified from single-cell transcriptomic data. (C) Stacked bar chart showing changes in the proportion of OPC subtypes at P30, P180, P360 and P720. (D, E) UMAP plots (D) and a heatmap (E) showing expression levels of *Vim, Dbn1,* and *Flnb* across OPC subtype at different ages. *P*-values were calculated using one-way ANOVA test.

To achieve greater cellular resolution, we interrogated these genes across functionally defined OPC subtypes using single-cell transcriptomic data spanning P30 to P720 [62]. These data reflected an age-dependent shift in OPC heterogeneity, characterized by a decrease in proliferative and differentiating populations alongside an expansion of the quiescent pool (Figure 6B, C). Within these subtypes, the expression of *Vim*, *Dbn1*, and *Flnb* varied distinctly. UMAP analysis demonstrated that *Flnb* levels remained low across all ages. In contrast, *Vim* expression declined with age in transitioning OPCs yet exhibited an increase in differentiating OLs (Figure 6D, E). Proteomic analysis confirmed that Vim and Dbn1 protein levels mirrored these transcriptional trends (Supplementary Figure S7A) [5].

Additionally, qRT-PCR showed that *Vim* mRNA was reduced in OLs, whereas *Dbn1* and *Flnb* were elevated in OLs (Supplementary Figure S7B). Specifically, *Vim* expression was found to be downregulated to approximately 0.2-fold in OLs compared to OPCs, while *Dbn1* levels increased to approximately 2.5-fold and *Flnb* showed a massive induction of over 120-fold relative to OPCs. Due to its dynamic regulation during differentiation, significant dysregulation in multiple sclerosis lesions, and high sensitivity to aging, we focused our further functional characterization on *Vim*.

### O-GlcNAcylation of Vimentin is critical for OPC differentiation

Our proteomic analysis identified two O-GlcNAcylation sites on Vimentin, Threonine 35 (T35) and Threonine 63 (T63) (Supplementary Figure S8). To assess whether these modifications alter the intrinsic properties of Vimentin, we exogenously expressed HA (N-terminal)-tagged wild-type (*Vim*^WT^) or an O-GlcNAc-deficient mutant (*Vim*^T35A^ ^T63A^) in HEK293T cells. Immunostaining confirmed that both variants exhibited similar subcellular localization (Figure 7A, B).

**Figure 7.**
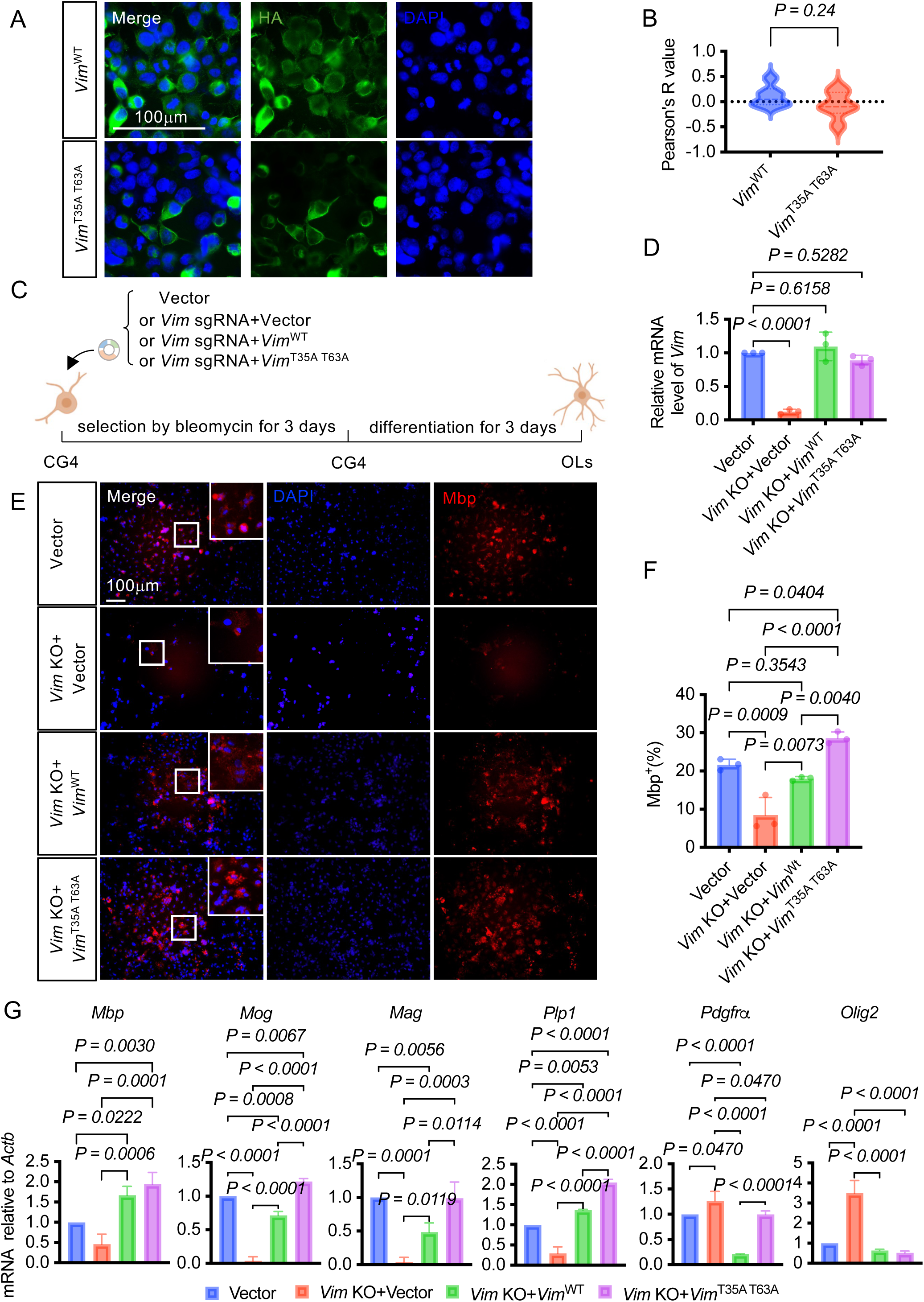
Functional validation of Vimentin O-GlcNAcylation in oligodendrocyte differentiation. (A, B) Localization of HA-tagged *Vim*^WT^ and *Vim*^T35AT63A^ in HEK293T cells. Scale bar: 100 μm. n=6 independent biological experiments. (C) Schematic diagram of the experimental workflow. CG4 cells were transfected with constructs for *Vim* KO, *Vim*^WT^ rescue, or *Vim*^T35A^ ^T63A^ rescue. Subsequently, the transfected cells were subjected to bleomycin selection for 3 days, during which OPC differentiation was simultaneously induced. (D) qRT-PCR analysis to evaluate the efficiency of *Vim* KO. The results showed that the expression level of *Vim* was reduced by approximately 90% in the *Vim* KO group compared with the control group. P-values were calculated using two-tailed unpaired Student’s t-test. (E) Representative images of CG4 cells at the stage of OL. Scale bar: 100 μm. n=3 independent biological experiments. (F) Statistical analysis of the positive rate of mature OLs (Mbp^+^ cells) in CG4 cells. P-values were calculated using one-way ANOVA test. (G) qRT-PCR analysis to detect the mRNA expression levels of OL maturation-related genes (*Mbp, Mog, Mag,* and *Plp1*) and OPC related genes (*Pdgfr*α and *Olig2*). P-values were calculated using one-way ANOVA test. n=3 independent biological experiments.

To investigate the role of Vimentin O-GlcNAcylation in OPC differentiation, we utilized CRISPR-Cas9 technology to knock out *Vimentin* in the CG4 OPC line, followed by rescue with either *Vim*^WT^ or the O-GlcNAc-deficient mutant. Cells were selected with Bleomycin for 3 days and subsequently induced to differentiate for an additional 3 days (Figure 7C). Knockout efficiency was confirmed by qRT-PCR, revealing a ∼90% reduction in *Vim* mRNA (Figure 7D). Functional assays revealed that *Vim* knockout (*Vim* KO) impaired OPC differentiation, reducing the percentage of mature Mbp^+^ cells to 8.48% compared to 21.68% in vector controls. While re-expression of *Vim*^WT^ rescued this differentiation defect (18.04% Mbp^+^ cells), the O-GlcNAc-deficient *Vim*^T35A^ ^T63A^ mutant exhibited a significant gain-of-function effect, further enhancing maturation to 28.59% Mbp^+^ cells (Figure 7E, F). In addition, we performed qRT-PCR to evaluate the expression of a panel of mature oligodendrocyte (*Mbp, Mog, Mag,* and *Plp1*) and OPC-related (*Pdgfr*α and *Olig2*) genes (Figure 7G). Consistent with the immunostaining data, the expression of all these oligodendrocyte genes was downregulated to less than 0.5-fold of the vector control levels in *Vim* KO cells compared to controls. This defect was rescued by *Vim*^WT^ while the *Vim*^T35A^ ^T63A^ enhanced rescue efficiency, increasing the expression of *Mbp* to approximately 2.0-fold, *Mog* to approximately 1.2-fold and *Plp1* to approximately 2.0-fold relative to the vector control. Regarding OPC-related genes, *Vim* KO led to aberrantly elevated *Olig2* and *Pdgfr*α levels to approximately 3.5-fold and 1.2-fold compared to the vector control respectively, which were effectively restored to control levels by both *Vim*^WT^ and *Vim*^T35A^ ^T63A^ rescues.

Furthermore, we performed Ki67 and TUNEL staining (Supplementary Figure S9A, C). The results showed no significant differences in the proportion of Ki67^+^ or TUNEL^+^ cells between Vector control, *Vim* KO, and rescue groups (Supplementary Figure S9B, D).

Finally, to validate the identified O-GlcNAcylation sites of Vimentin, we performed immunoprecipitation assays. We expressed HA (C-terminal)- and Flag (N-terminal)-tagged wild-type (Vim^WT^) or mutant (Vim^T35A^ ^T63A^) Vimentin in CG4 cells and enriched the proteins using anti-Flag beads. Western blot analysis revealed that, though not completely abolished, the O-GlcNAcylation level of the Vim^T35A^ ^T63A^ mutant was reduced to approximately 50% compared to Vim^WT^ (Figure 8A-C). To rule out the possibility of protein degradation, we quantified the ratio of the C-terminal HA tag to the N-terminal Flag tag (Figure 8D). The HA/Flag ratio remained comparable between Vim^WT^ and mutant Vim^T35A^ ^T63A^, indicating that the protein stability and full-length integrity were maintained. These results support that the observed reduction in O-GlcNAc signal is largely due to the loss of the modification sites at Threonine 35 and Threonine 63.

**Figure 8.**
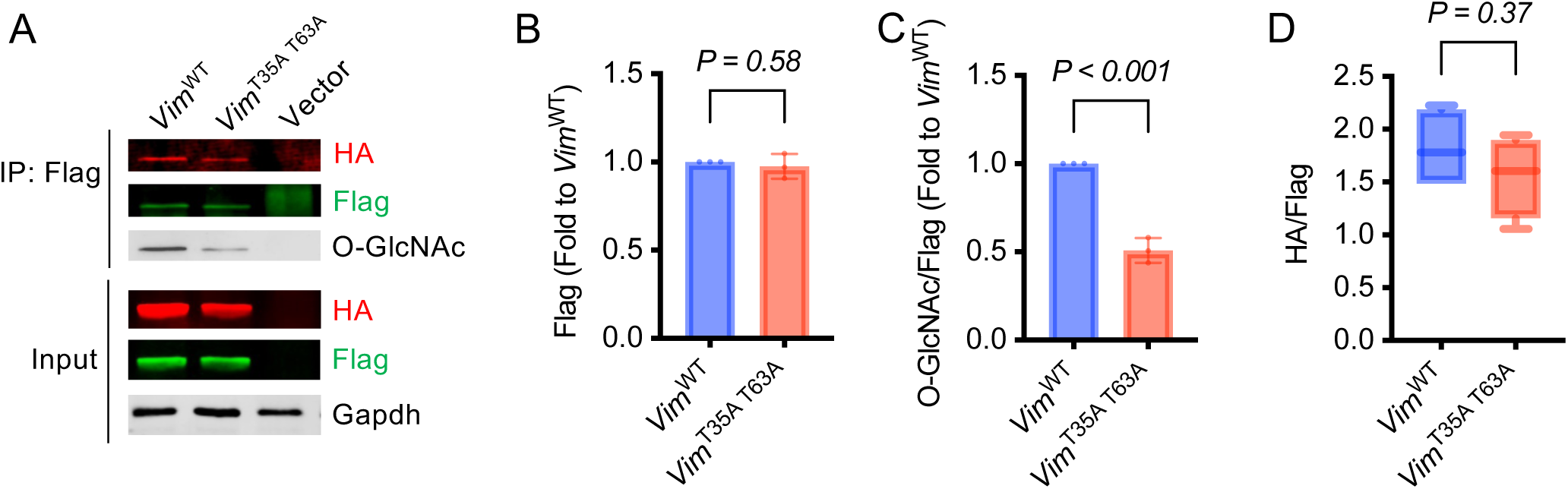
Analysis of Flag-HA-tagged Vimentin O-GlcNAcylation validation. (A) Representative results of IP experiments comparing O-GlcNAcylation levels between Vim^WT^ and Vim^T35A^ ^T63A^ and WB of Flag-HA-tagged Vim^WT^ and Vim^T35A^ ^T63A^. Samples were collected from CG4. (B-D) Statistical analysis of the relative Flag levels (B), O-GlcNAc levels normalized to Flag (C) and HA normalized to Flag (D). n=3 independent biological experiments. P-values were calculated using two-tailed unpaired Student’s t-test.

## Discussion

In this study, we profiled the O-GlcNAcome in neonatal mouse OPCs, identifying 165 O-GlcNAcylation sites across 118 proteins, predominantly nuclear and monosubstituted. Comparative analysis revealed 74 novel sites on 67 proteins, including 22 uniquely O-GlcNAylated in OPCs, enriched in intrinsically disordered regions, linear interaction motifs, and pathways related to cytoskeletal organization and CNS development. Molecular docking simulations indicated intermediate binding affinities to O-GlcNAc transferase (Ogt). Integration with transcriptomic datasets from multiple sclerosis and aging contexts identified candidate proteins, including *Vimentin (Vim*). Functional validation confirmed Thr35 and Thr63 as O-GlcNAc sites on Vimentin, demonstrating that inhibiting this modification enhances OPC differentiation. This work establishes the first comprehensive O-GlcNAcome atlas for OPCs, providing an essential resource for elucidating myelination mechanisms and their dysregulation in demyelinating disorders.

Consistent with prior observations that OGT preferentially targets IDRs[34, 57], our data reinforce that O-GlcNAcylation is enriched in these conformationally flexible, non-folded domains (69.7% of sites). The flexible, non-folded structure of IDRs is essential for dynamic protein-protein interactions, molecular recognition, and cellular signaling regulation [58]. Because O-GlcNAcylation often occurs within these disordered regions, it is uniquely positioned to modulate the conformational plasticity of its targets, thereby regulating their biological functions. This structural modulation provides a biochemical basis for O-GlcNAcylation to act as a functional bridge between molecular signaling and complex neurodevelopmental processes [26, 27, 63]. LIPs, a functional subset of IDRs with intrinsic binding capability[64], were also heavily modified in our dataset (35.2%). Site-specific O-GlcNAcylation within LIPs is therefore likely to exert potent effects on rewiring protein interaction networks that govern OPC process extension, migration, and terminal differentiation.

Although global O-GlcNAcylation fluctuates with age, underlying site-specific alterations are unresolved. Consequently, it is imperative for future studies to employ high-resolution, quantitative O-GlcNAcome profiling to capture the nuances of temporal differentiation. Currently, the technical challenges in direct detection of Vimentin O-GlcNAcylation in mature OLs preclude a precise stoichiometric comparison across developmental stages. Our observation of increased Mbp^+^ cell density likely reflects a shift in lineage commitment, suggesting potential alterations in the efficiency of oligodendrogenesis. Further mechanistic investigations should prioritize the identification of additional molecular candidates and the functional crosstalk between PTMs, particularly the O-GlcNAcylation-phosphorylation axis. These endeavors hold the potential to reveal novel therapeutic targets for alleviating differentiation arrest and fostering remyelination in demyelinating pathologies.

## Acknowledgement

We thank Mingjie Chen (Shanghai NewCore Biotechnology Co., Ltd.) for providing data analysis and visualization support.

## Funding

This work was supported by grants from the National Natural Science Foundation of China (32370853).

## Data availability

The O-GlcNAcome data have been deposited in PRIDE (Project PXD043506, https://www.ebi.ac.uk/pride/). Transcriptomic data about mouse OPCs across lifespan have been deposited in the Sequence Read Archive (SRA, https://www.ncbi.nlm.nih.gov/) under SUB15247167.

## Notes

### Competing Interest Statement

The authors have declared no competing interest.

