## Supplementary Figure 1-9 for "Global O-GlcNAcome Identifies O-GlcNAcylated Proteins Regulating Oligodendrocyte Precursor Cells Differentiation"

**Supplementary Data**

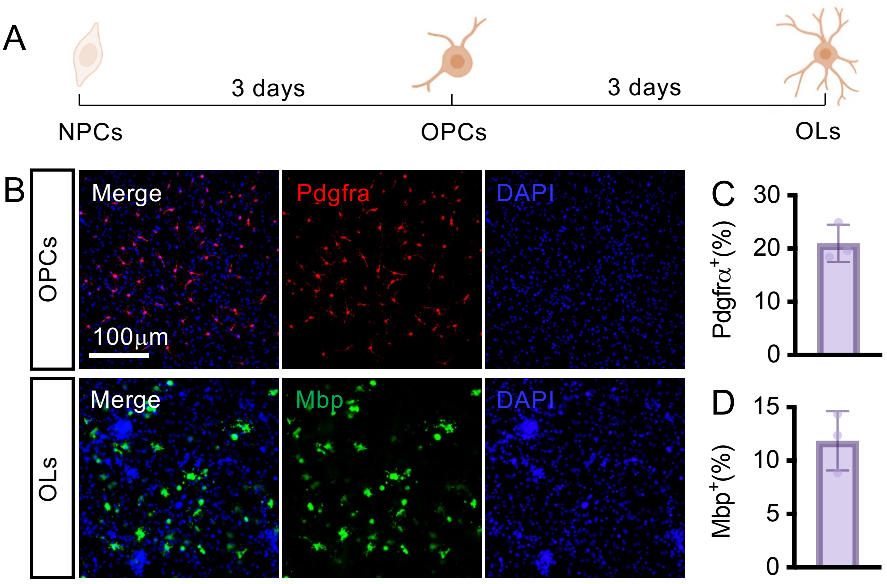


**Supplementary Figure S1. Culture purity of OPC and mature OL differentiated from NPC**

(A) Schematic timeline of NPC-to-OL differentiation. NPCs were first induced to differentiate into OPCs (culture duration: ~3 days), and OPC differentiation efficiency was evaluated at this stage; subsequently, OPCs were further differentiated into mature OLs (additional culture duration: ~3 days), with OL differentiation efficiency assessed at the final stage. (B) Representative image of NPC-derived OPC (Pdgfrα$^{+}$) and OL (Mbp$^{+}$). n=3 independent biological experiments. Scale bar: 100 μm. (C, D) Quantitative analysis of culture purity for NPC-derived OPCs (C) and mature OLs (D). Purity was determined by calculating the percentage of Pdgfrα$^{+}$ cells (for OPCs) or Mbp$^{+}$ cells (for OLs) relative to the total number of DAPI-stained nuclei.

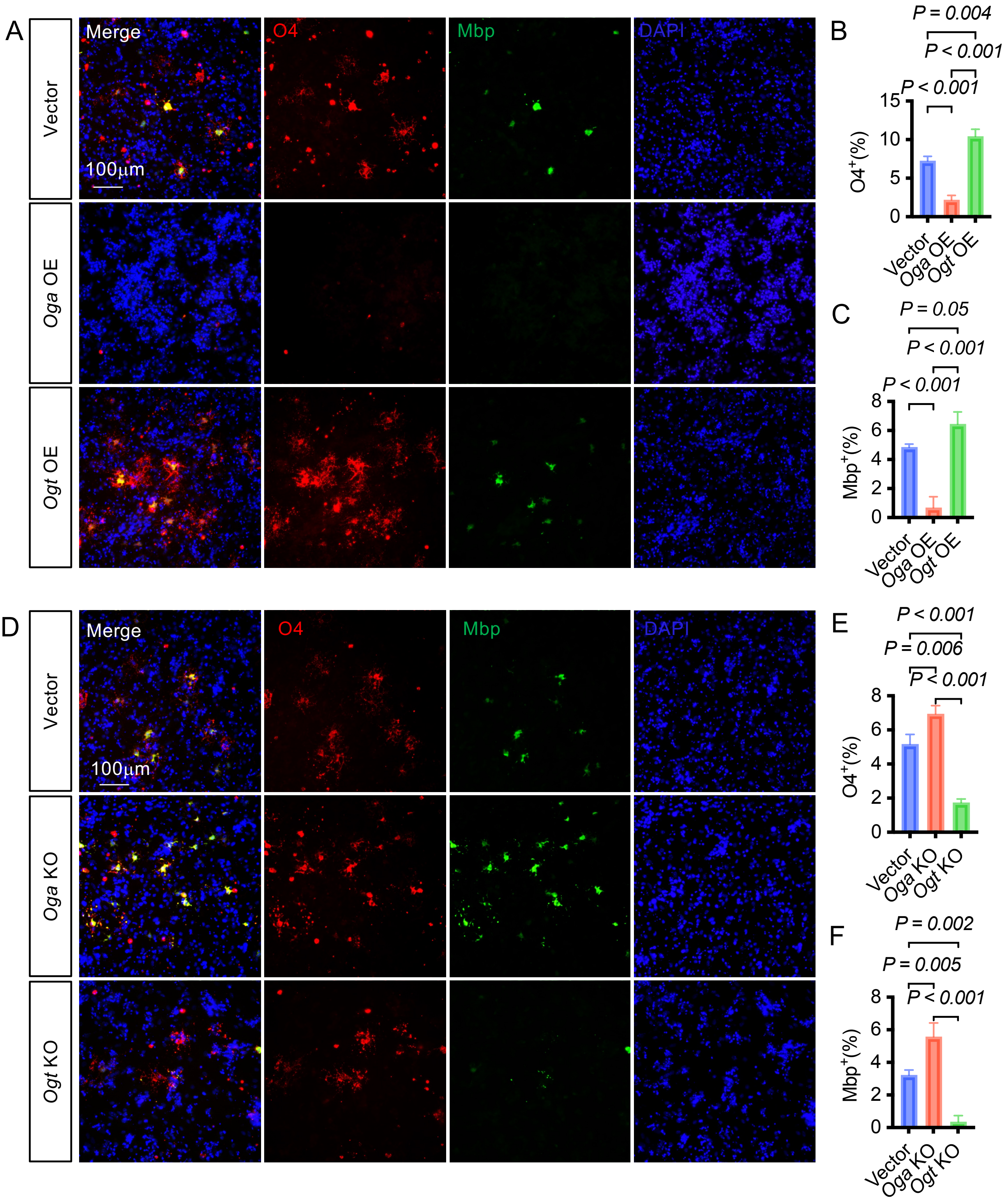


**Supplementary Figure S2. Overexpression and Knockout of *Oga* and *Ogt* modulates the differentiation of OPCs into mature OLs**

(A) Representative images showing the effect of *Oga* and *Ogt* overexpression on the differentiation potential of OPCs into OLs ($O4^+$) and mature OLs ($Mbp^+$). Scale bar: 100 μm. (B, C) The proportion of OL ($O4^+$ and $Mbp^+$) of *Oga* and *Ogt* overexpression. Quantitative analysis of the percentage of $O4^+$ (B) and $Mbp^+$ (C) OLs in OPC cultures transfected with *Oga* and *Ogt* overexpressing plasmids versus empty vector controls. (D) Representative images showing the effect of *Oga* and *Ogt* knockout on the differentiation potential of OPCs into OLs ($O4^+$) and mature OLs ($Mbp^+$). Scale bar: 100 μm. (E, F) The proportion of OL ($O4^+$ and $Mbp^+$) of *Oga* and *Ogt* knockout. Quantitative analysis of the percentage of $O4^+$ OLs (E) and $Mbp^+$ OLs (F) in OPC cultures transfected with *Oga* and *Ogt* overexpressing plasmids versus empty vector controls. n=3 independent biological experiments. P-values were calculated using one-way ANOVA test. n=3 independent biological experiments.

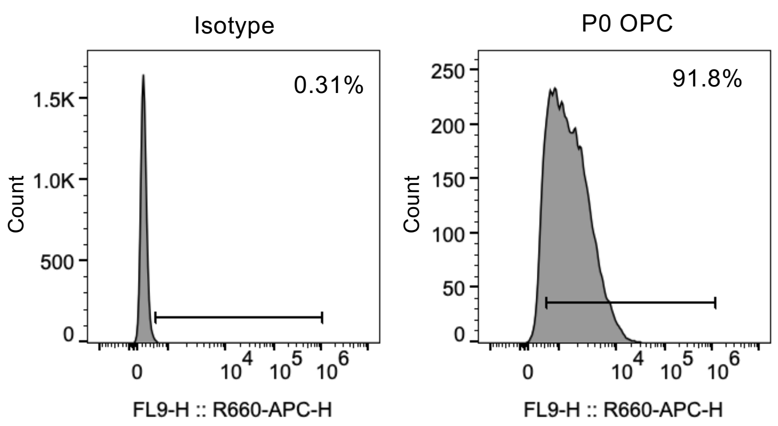


**Supplementary Figure S3. Purity of OPC used for LC-MS**

Flow cytometry was performed to determine the purity of primary OPCs isolated P0 mice, using an antibody against Pdgfrα (Invitrogen). An isotype control antibody was used to establish the background signal and gate the positive population. Representative flow cytometry plots showed that the percentage of Pdgfrα$^{+}$ cells in the isotype control group was 0.31% (indicating minimal non-specific staining), while the percentage of Pdgfrα$^{+}$ OPCs in the P0 OPC preparation was 91.8%. This high Pdgfrα positive rate confirms that the OPCs used for subsequent LC-MS analysis had sufficient purity, ensuring the reliability and specificity of the downstream proteomic or metabolomic data generated from these cells.

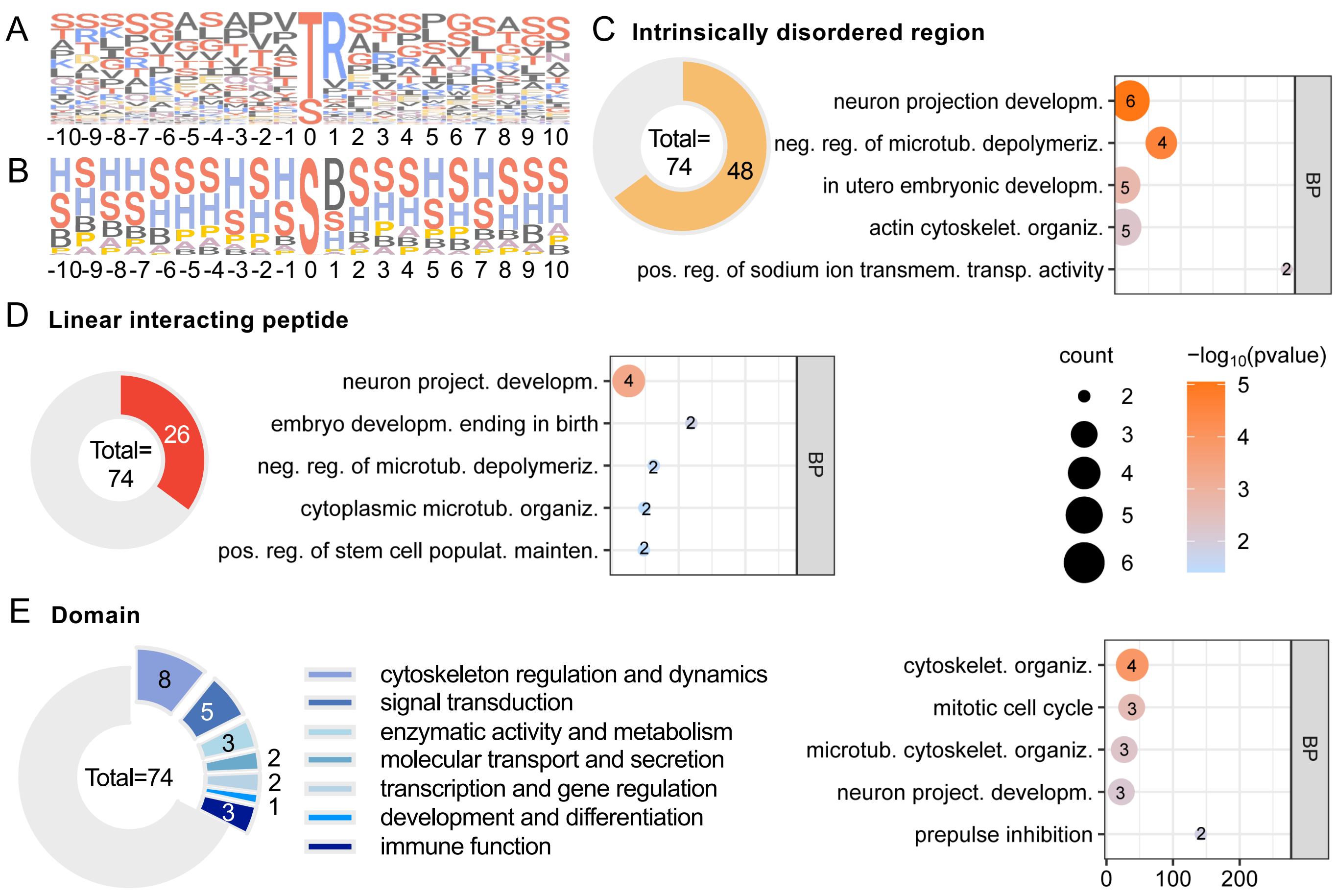


**Supplementary Figure S4. Motif and structural analysis of OPC-specific O-GlcNAcylated sites**
(A, B) Motif analysis of the amino acid sequences (A) and the chemical properties of amino acid (B) surrounding the OPC-specific O-GlcNAcylated sites. (C) Distribution of O-GlcNAcylated sites within intrinsically disordered regions (IDRs) and GO enrichment analysis of the corresponding proteins. (D) Distribution of O-GlcNAcylated sites within linear interaction peptides (LIPs) and GO enrichment analysis of the corresponding proteins. (E) Distribution of O-GlcNAcylated sites within structured protein domains and the functional classification of the corresponding proteins.

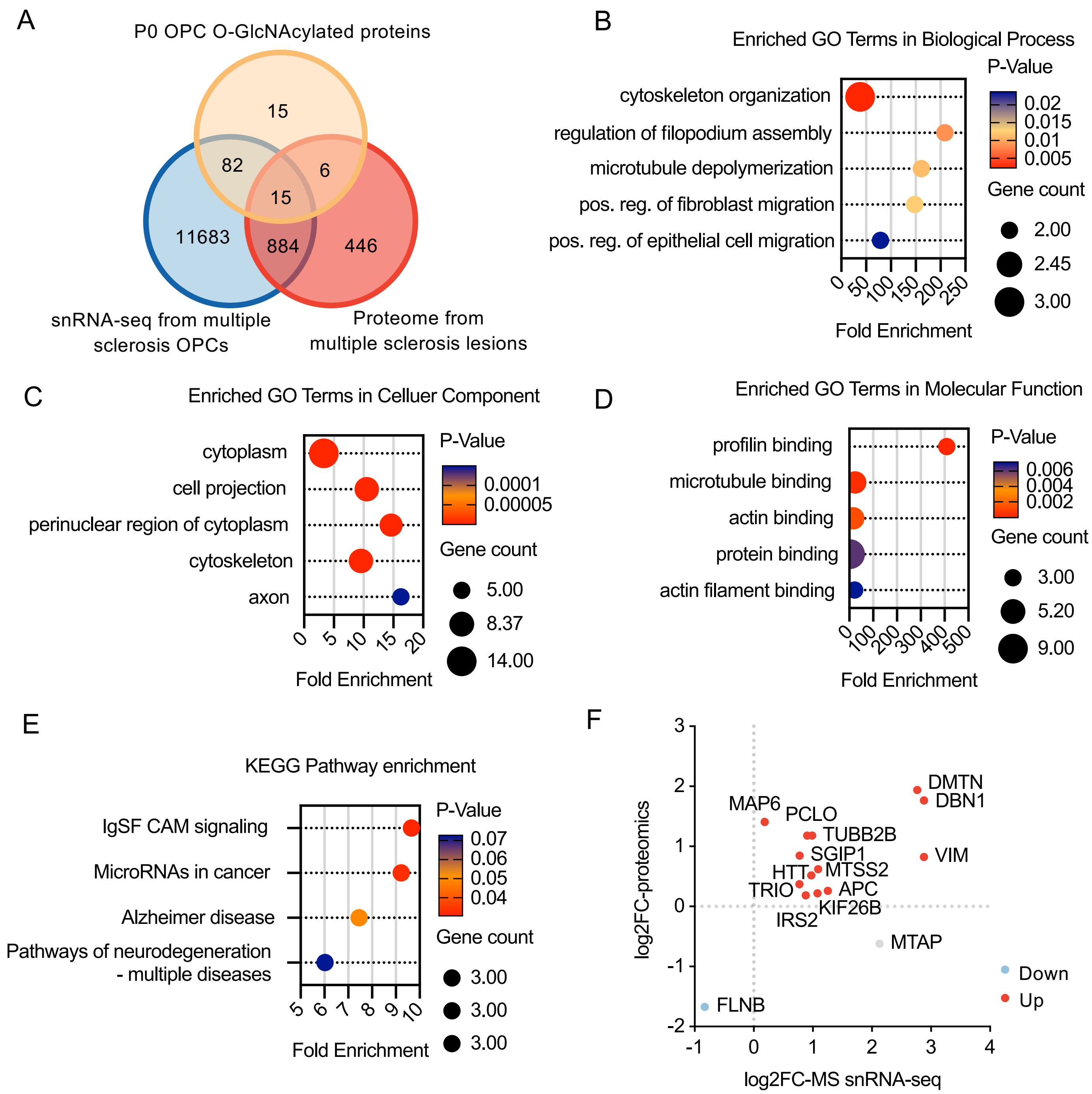


**Supplementary Figure S5. Association of OPC O-GlcNAcylated proteins with multiple sclerosis** (A) Venn diagram showing 15 overlapping proteins among the OPC O-GlcNAcylated proteins, single-nucleus RNA sequencing (snRNA-seq) data from people with multiple sclerosis, and proteomic data from multiple sclerosis lesions. (B-D) Bubble chart of enriched GO terms in biological process (B), cellular component (C) and molecular function (D) for OPC O-GlcNAcylated proteins with multiple sclerosis. (E) KEGG pathway enrichment analysis for OPC O-GlcNAcylated proteins with multiple sclerosis. (F) Scatter plot showing the expression changes (log2 fold change) of the 15 overlapping proteins in multiple sclerosis snRNA-seq (x-axis) and multiple sclerosis lesion proteomics (y-axis) data.

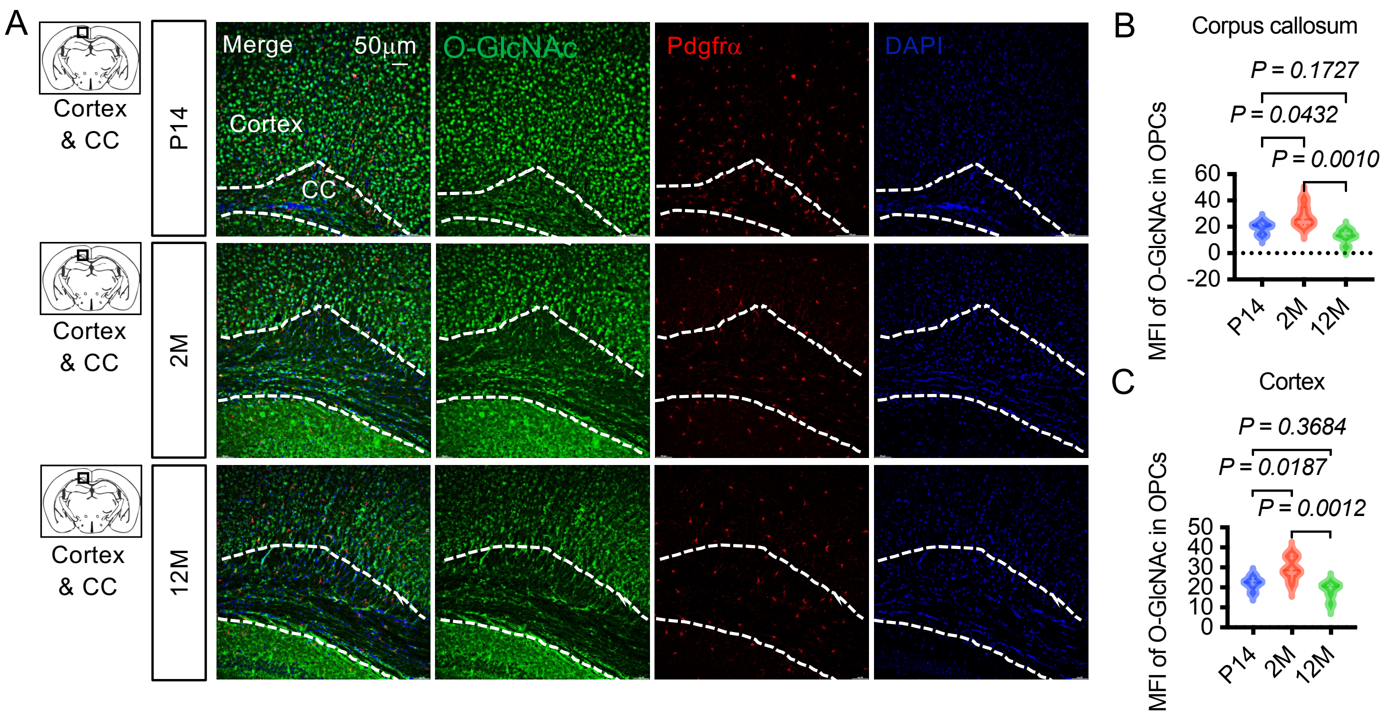


**Supplementary Figure S6. O-GlcNAc level of OPC of different ages**

(A) O-GlcNAc level of OPC (Pdgfra$^{+}$) at P14, 2M, 12M. Scare bar: 50µm. (B, C) Statistical analysis of the O-GlcNAc levels corresponding to OPC in the corpus callosum (CC) (B) and cortex (C). n=6 independent biological experiments. P-values were calculated using one-way ANOVA test.

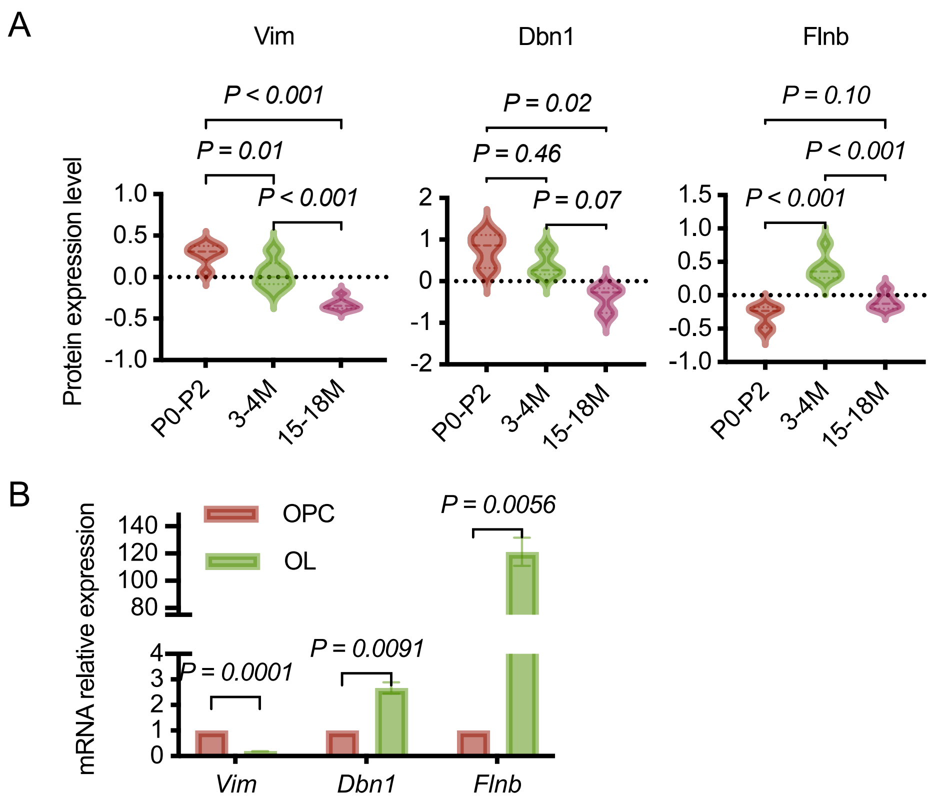


**Supplementary Figure S7. Expression of *Vim*, *Dbn1*, and *Flnb* across age and cell types**

(A) Proteomic data showing age-related changes in Vim and Dbn1 and Flnb protein abundance in OPCs (P0-P2, 3-4M, 15-18M). (B) qPCR revealing high *Vim* mRNA expression in OPCs, significantly reduced in OLs, with *Dbn1* and *Flnb* showing elevated expression in OLs. Data are presented as mean ± SD. P-values were calculated using two-tailed unpaired Student's t-test and one-way ANOVA test.

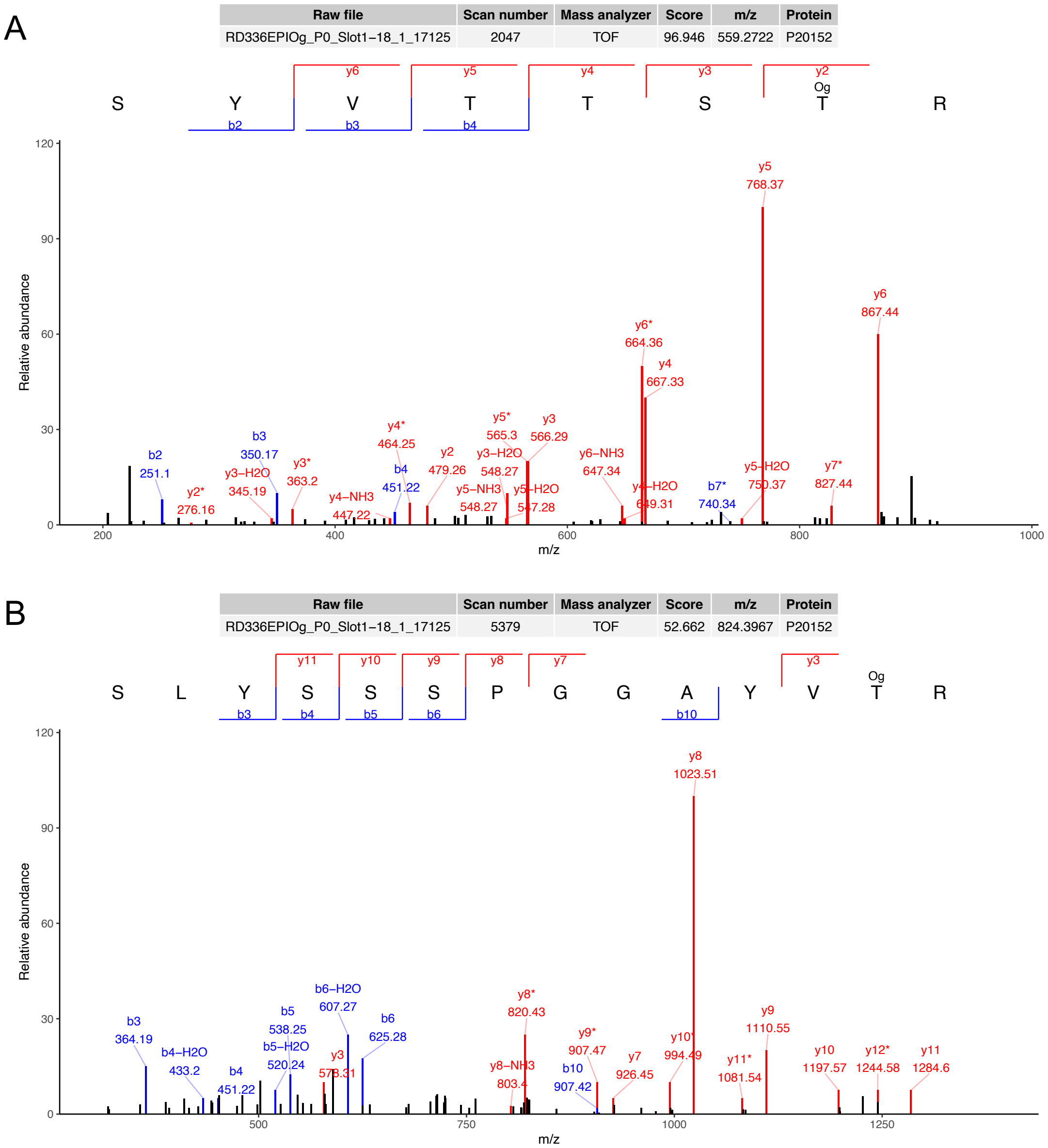


**Supplementary Figure S8. MS/MS spectra for the O-GlcNAcylated peptides of Vimentin**

(A, B) MS/MS spectra validating site-specific O-GlcNAcylation of Vimentin at Threonine 35 (T35) (A) and Threonine 63 (T63) (B), isolated from P0 mouse primary OPCs.

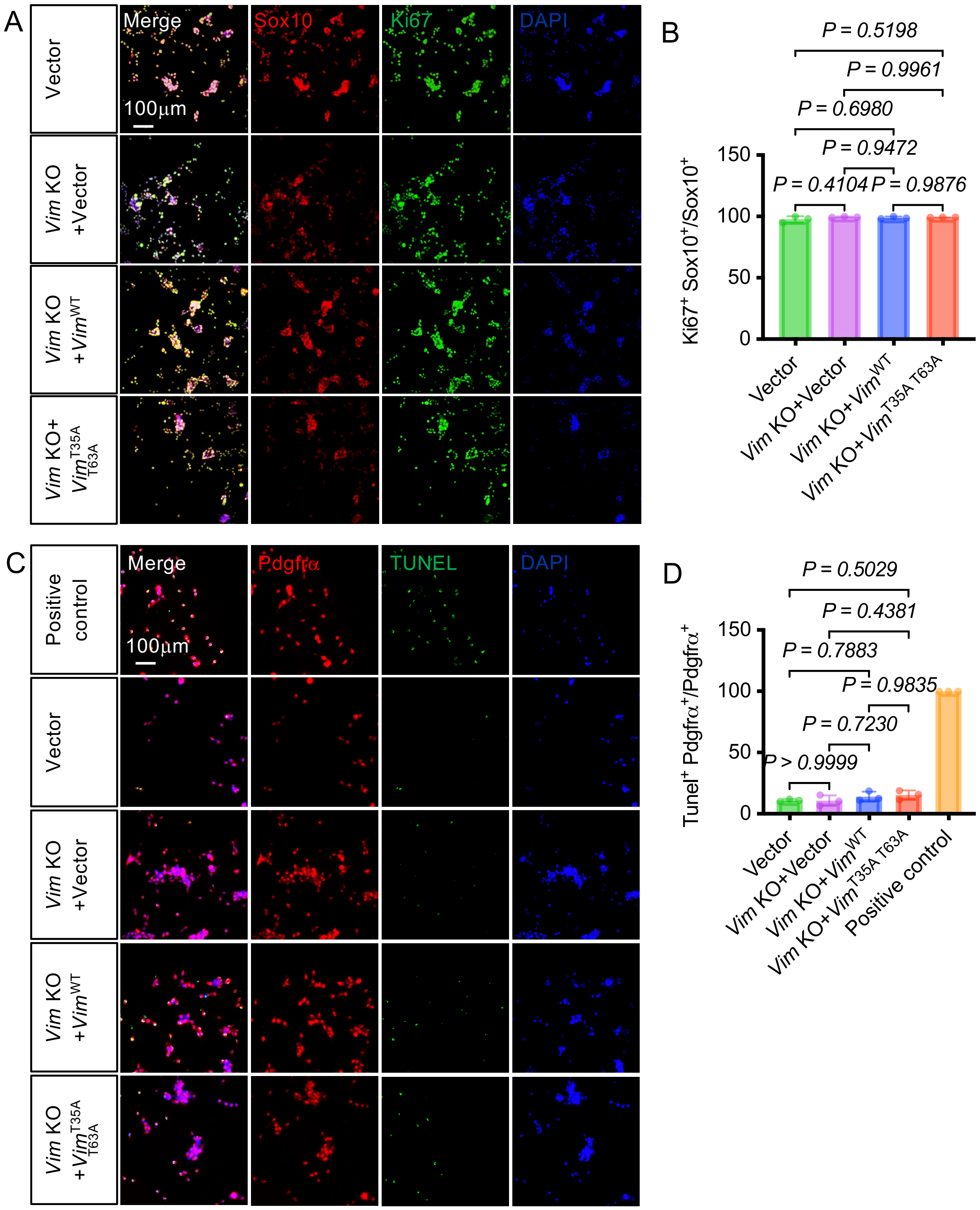


**Supplementary Figure S9. Influence of *Vimentin* on the proliferation and apoptosis of OPCs**

(A) Representative images of Sox10 and Ki67 co-staining in CG4 cells under empty vector control, *Vim* KO+Vector, and *Vim* rescue conditions. Scale bar: 100 μm. (B) Quantitative analysis of the ratio of $Sox10^{+}Ki67^{+}$ cells to total $Sox10^{+}$ cells. Statistical results demonstrate that neither *Vim* KO nor *Vim* rescue exerts a significant effect on the proliferation of CG4 cells. (C) Representative immunofluorescence images of Pdgfrα and TUNEL co-staining in CG4 cells under vector control, *Vim* KO+Vector, and *Vim* rescue conditions. Scale bar: 100 μm. (D) Quantitative analysis of the ratio of $Pdgfr\alpha^{+}TUNEL^{+}$ cells to total $Pdgfr\alpha^{+}$ cells. Statistical results indicate that *Vim* knockout and *Vim* rescue do not have a notable impact on the apoptosis of CG4 cells. P-values were calculated using one-way ANOVA test. n=3 independent biological experiments.
